# Cooperativity, dynamics, and the free-energy surfaces of charge-patterned IDPs

**DOI:** 10.64898/2026.05.21.726897

**Authors:** Valentin von Roten, Miloš T. Ivanović, Soundhararajan Gopi, Andrea Holla, Andreas Prestel, Mark Nüesch, Ketty C. Tamburrini, Daniel Nettels, Birthe B. Kragelund, Robert B. Best, Benjamin Schuler

## Abstract

The free-energy surfaces that underlie the conformational distributions of intrinsically disordered proteins (IDPs) are shallow and lack the deep minima characteristic of stable, folded structures. However, even in the absence of secondary or tertiary structure, sequence patterning can lead to conformational preferences and changes in chain dimensions as a function of solution conditions. While patterning effects have received extensive attention from simulation and theory, there is little corresponding data from experiment. Here we investigate the impact of charge patterning on chain dimensions and dynamics in a set of specifically designed polyampholytic IDP variants across the natural range of charge segregation with single-molecule FRET, nanosecond fluorescence correlation, circular dichroism, and NMR spectroscopy. We find that the conformational ensembles and their cooperative response to salt concentration show prominent and systematic dependencies on charge patterning, and to some extent on residue type. In contrast, the chain dynamics remain in the tens-of-nanosecond range, consistent with the absence of pronounced free-energy barriers. In close combination with molecular simulations, we show how the concept of susceptibility can be used to quantify cooperativity in the absence of barriers and relate it to the shallow free-energy surfaces of IDPs.

## Introduction

Many proteins do not fold into a well-defined 3D structure or contain large disordered regions, and thus sample large conformational ensembles^1-3^. These intrinsically disordered proteins (IDPs) are particularly prevalent in signaling pathways and cellular regulation, and they can enable rapid binding kinetics and higher-order complex formation^3,4^. In contrast to the classical view that protein-protein interactions require structurally well-defined complementary interaction surfaces, IDRs can remain disordered even in their bound state^5-11^. The amino acid sequence has a pronounced influence on both the interactions within the chain — and thus the chain dimensions — as well as the interactions between IDPs^12-15^, which also determine their propensity to form biomolecular condensates^16-18^. IDPs are often polyampholytes, i.e., they are rich in both acidic and basic amino acid residues^13,15^, and the electrostatic interactions between them have a particularly large effect on the dimensions and conformations of IDPs^19-21^.

As a result of the absence of tertiary structure, IDPs lack the pronounced free-energy minima of folded proteins, i.e., their free-energy surfaces are comparatively shallow^22,23^. Correspondingly, concepts from polymer theory are often useful for describing and quantifying the conformational distributions of IDPs and their dependence on solution conditions^24-27^. However, even in the absence of structure formation, conformational preferences can emerge from sequence patterns in IDPs^28,29^. For polyampholytic IDPs, for instance, not only the total number of acidic and basic residues^30,31^ is of importance, but also their distribution within the sequence^12,15,30-32^, and both have been related to the chain dimensions of IDPs in solution^15,30-34^ as well as their dynamics^35,36^. However, systematic studies of these effects have largely been a focus of simulation and theory; experimental investigations are sparse, and quantitative assessments of the cooperativity of the conformational changes, the resulting free-energy surfaces, and the underlying chain dynamics are lacking.

Here, we designed a model system based on naturally occurring IDP sequences to systematically probe the influence of charge segregation and sequence composition with a combination of experimental techniques and molecular simulations: NMR and circular dichroism (CD) spectroscopy allow us to assess structural propensities; single-molecule Förster resonance energy transfer (FRET) to probe intrachain distance distributions; nanosecond fluorescence correlation spectroscopy (nsFCS) to quantify chain dynamics; and coarse-grained and all-atom explicit-solvent simulations benchmarked by the experimental data enable us to interpret the measurements in molecular detail.

## Results

We designed a molecular system that allows us to probe the influence of both charge segregation and sequence composition. As a starting point for polyampholyte sequences with high, yet realistic — i.e., naturally occurring — charge density, we selected 29-residue segments of two highly charged IDPs, the basic human linker histone H1.0 (H1) and its nuclear chaperone, the acidic prothymosin α (ProTα)^5,8,38^. By fusing the two segments, we obtained a diblock-copolymeric sequence with a total length well suited for probing its end-to-end distance distribution and dynamics with single-molecule FRET^39^, and including terminal Cys residues for fluorophore labeling^40^ (Fig. 1a). In this sequence, 14 positively charged Lys residues are present in the N-terminal half, and 14 negatively charged residues — 12 Glu and 2 Asp — in the C-terminal half. We refer to this variant as KE_high_, indicating the predominant types of charged residues and the high degree of charge segregation^28,37^ (Fig. 1c). The level of charge segregation of KE_high_ is comparable to values at the extreme end of the distribution for polyampholytic IDPs naturally occurring in the human proteome (Fig. 1d). To vary the degree of charge segregation while keeping the sequence composition identical, we first swapped the two central 15-residue segments, resulting in a tetrablock-copolymeric variant with four alternating charge clusters, KE_mid_; in a second step, we generated KE_low_, an octablock variant with eight alternating charge clusters, corresponding to the lowest degree of charge segregation we investigated. The resulting sequences thus provide samples across the natural distribution of charge segregation in IDPs (Fig. 1d). To additionally probe the influence of the chemical identity of the charged residues, we created variants with sequence and charge patterning identical to KE_low_, but with Lys residues replaced by Arg (RE_low_), Glu residues replaced by Asp (KD_low_), or both (RD_low_) (Fig. 1c; the five most C-terminal Lys residues were not replaced by Arg to reduce attractive interactions with the dyes^31^). These six variants were expressed recombinantly and purified.

**Figure 1.**
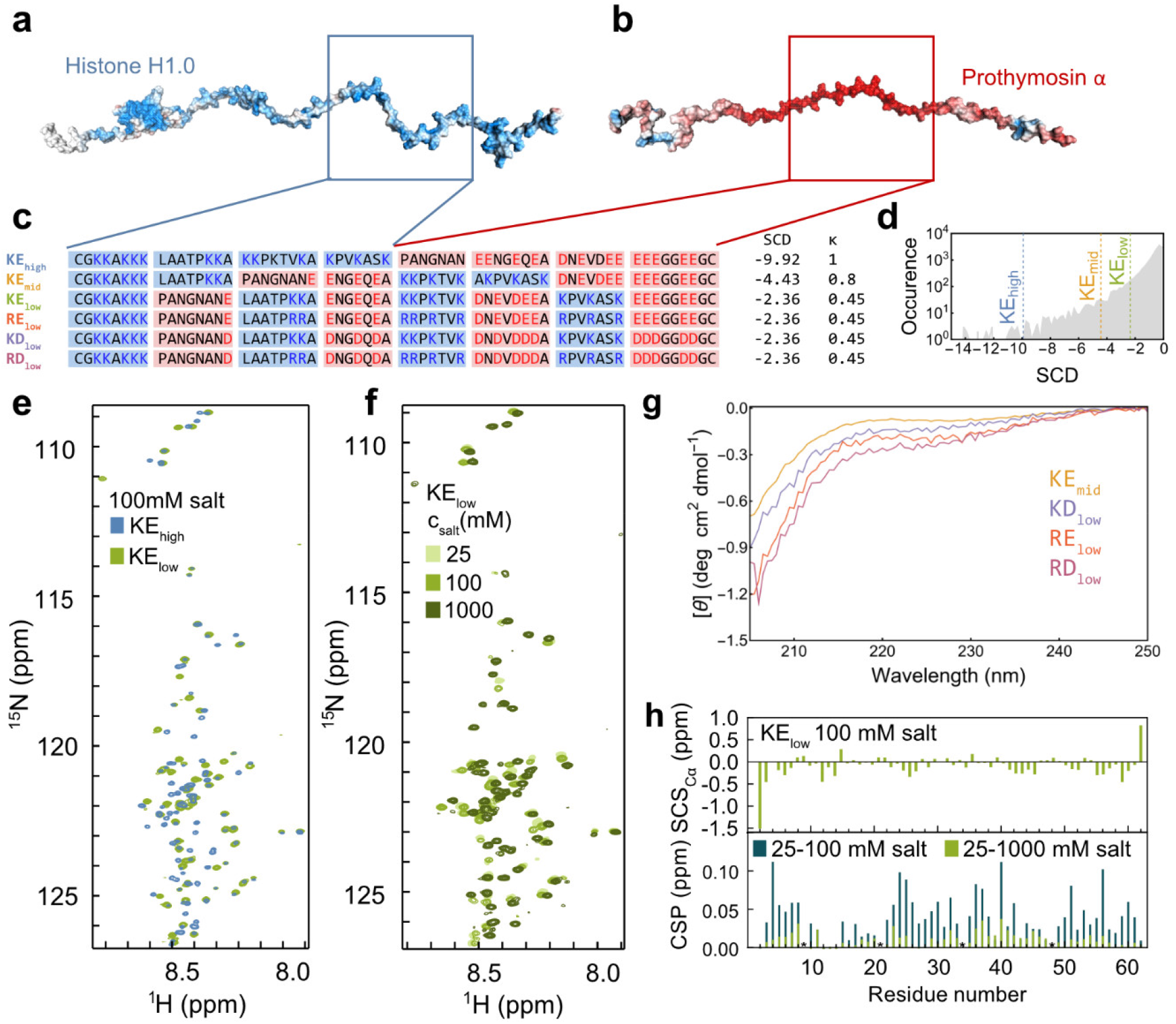
Disordered polyampholytic sequences designed for probing the effects of charge segregation and residue type. **a, b**, Segments of human histone H1.0 (H1) (**a**) and prothymosin α (ProTα)^5^ (**b**) were combined to generate the polyampholytic sequence KE_high_ (positively charged residues in blue, negatively charged in red). **c**, Sequences resulting from this combination, charge shuffling, and changes in amino acid type, with their acronyms, amino acid sequences, sequence charge decoration (SCD)^37^, and κ values^28^. **d**, Distribution of SCD for the disordered sequences of the human proteome, showing only the negative-SCD regime of polyampholytes (see Methods), with the values for KE_high_, KE_mid_, KE_low_ indicated as vertical dashed lines. **e**, ^1^H,^15^N-HSQC spectra of KE_high_ and KE_low_ at 100 mM salt. **f**, ^1^H,^15^N-HSQC spectra of KE_low_ at different KCl concentrations (see legend). **g**, Far-UV circular dichroism spectra of several variants at 20 mM salt. **h**, Secondary chemical shifts (SCS_Cα_) of KE_low_ at 100 mM KCl (upper panel) and salt-induced chemical shift perturbations (CSP) of KE_low_ (lower panel) comparing 25 and 100 mM (light green) or 25 and 1000 mM KCl (dark green). Asterisks indicate proline residues.

The segments of H1 and ProTα we fused are disordered in the parent proteins^41-43^, even when bound to each other^5,44^; nevertheless, we tested for the possible presence of secondary or tertiary structure in our fusion variants by CD and NMR spectroscopy (Fig. 1e-h). Even at low ionic strength (20 mM KCl), the CD spectra exhibit the typical random-coil signature^45^ (Fig. 1g), and NMR data for the selected cases of KE_high_ and KE_low_ show the low dispersion of ^1^H chemical shifts in ^1^H,^15^N heteronuclear single quantum coherence (HSQC) spectra expected for disordered proteins^46^ (Fig. 1e). This lack of secondary or tertiary structure is further supported by the random coil-like backbone ^13^C^α^ secondary chemical shifts (SCS) of KE_low_ (Fig. 1h). Increasing the KCl concentration leads to upfield shifts of the resonances but no change in overall peak dispersion (Fig. 1f,h), indicating the absence of structure formation even at high salt concentration^47,48^. Similar salt-induced changes in NMR signals were observed for ProTα and the random coil peptide QQEDQQ (Fig. S1), further supporting this conclusion. We observed some loss of NMR peak intensity with increasing salt concentration, at least in part due to increasing radio frequency power dissipation in the sample, but altogether, the sequences are clearly disordered under the conditions we use.

### Effect of sequence properties on conformational ensembles and salt response

To assess the effect of electrostatic interactions on chain dimensions, we labeled the termini of all variants with donor and acceptor fluorophores for single-molecule FRET spectroscopy and measured the average transfer efficiency, ⟨*E*⟩, reporting on the average end-to-end distance, on freely diffusing molecules as a function of KCl concentration (Fig. 2a-c). At low salt concentrations, all variants exhibit similarly high transfer efficiencies above 0.9, indicating close average proximity of the chain ends, as expected from the attraction between opposite charges in such polyampholytes^25,28,37,49^. With increasing KCl concentration, we observe for all sequences a continuous shift of a single peak to lower transfer efficiency that saturates at ~1 M KCl, reflecting an expansion of the chains, qualitatively similar to the change in compactness of the complex between full-length H1 and ProTα as a function of salt concentration^50^. The similarity of this behavior for the two different FRET pairs indicates that the fluorophores have no major effect on the transition (Fig. S2). The relation of fluorescence lifetimes to mean FRET efficiencies shows that the chains rapidly sample broad interdye distance distributions (Fig. 2d, Fig. S3) and compact with increasing temperature (Fig. 2e), as expected for IDPs^24,45,51^. Most importantly, however, the charge patterning of the variants has a pronounced impact on the transition midpoint of their salt-induced expansion, ranging from 71 ± 14 mM for KE_low_ to 305 ± 31 mM for KE_high_ (Fig. 2b). Moreover, the steepness of the transitions at the midpoint increases with charge segregation. More charge-segregated sequences thus lead to stronger intrachain attraction and require higher salt concentrations to expand from compact to more open conformational ensembles. Consequently, the dimensions of the three sequence variants are very different at physiological salt concentrations (Fig. 2b).

**Figure 2.**
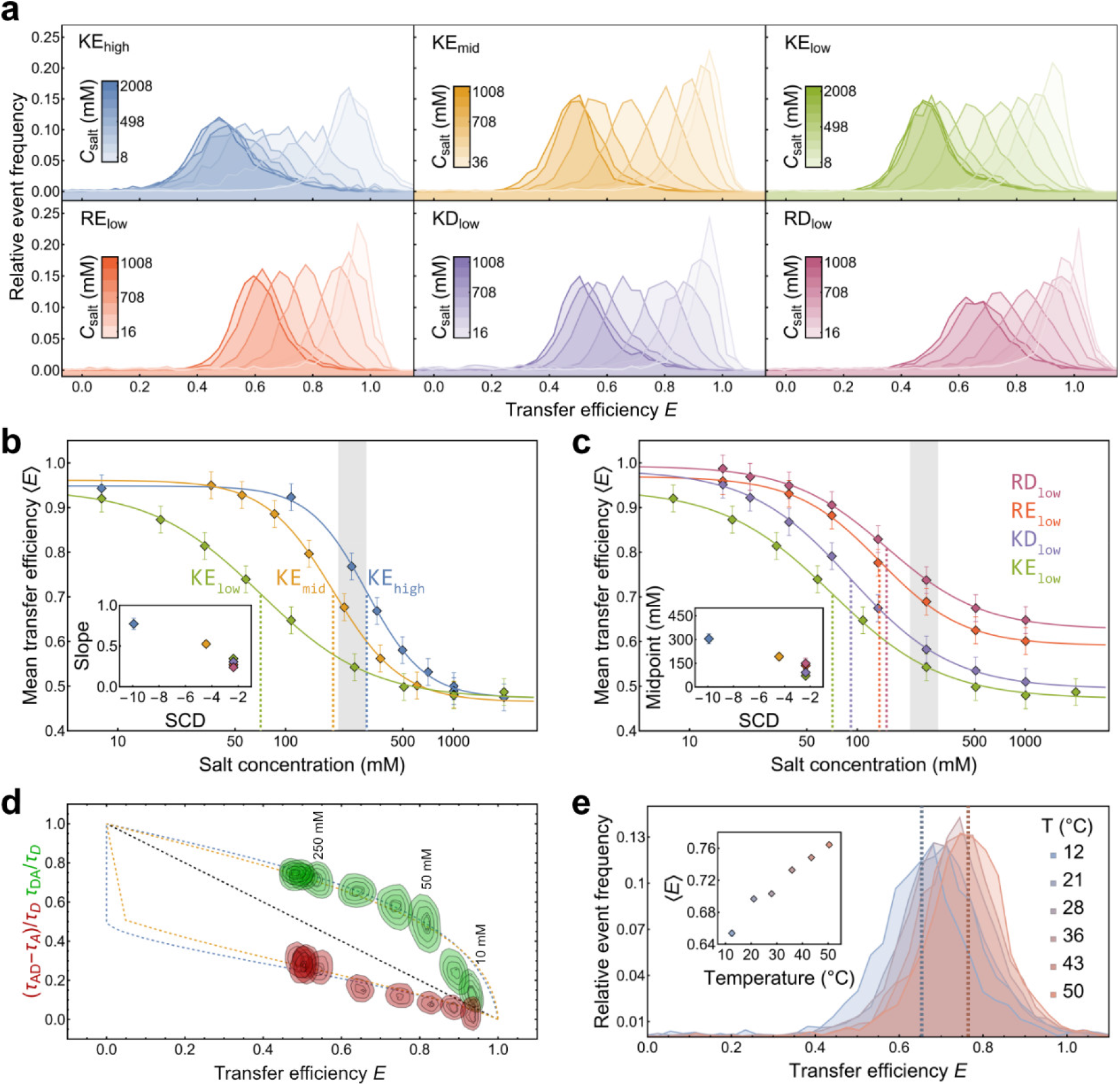
Impact of sequence and salt concentration on chain dimensions from single-molecule FRET experiments. **a**, KCl concentration dependence of transfer efficiency (E) histograms for the different sequence variants (color code as in Fig. 1c). **b**, Influence of charge segregation on salt concentration dependence of the mean transfer efficiency, ⟨E⟩. Error bars indicate the systematic uncertainty due to instrument calibration. Midpoints are highlighted by vertical dashed lines. The physiological salt concentration range is indicated as a gray band. Inset: slopes at the midpoints of the salt-induced transitions as a function of their SCD. **c**, Influence of amino acid type on salt concentration dependence of ⟨E⟩. Inset: transition midpoints for the different variants as a function of their SCD. Error bars indicate the systematic uncertainty due to instrument calibration. **d**, Transfer efficiency vs relative fluorescence lifetime for KE_high_ from single-molecule experiments at different salt concentrations (donor lifetime green, acceptor lifetime red). The black dashed line indicates the relation expected for static distances. The orange dashed lines show the relation for a rapidly sampled SAWv distance distribution, the blue dashed lines for a Gaussian chain. **e**, Temperature dependence of transfer efficiency histograms and mean transfer efficiency (inset) for KE_low_ at 100 mM salt.

Not only the degree of charge segregation influences the conformational properties, but also the type of charged residues: ⟨*E*⟩ as a function of salt concentration reveals interesting differences between KE_low_, RE_low_, KD_low_, and RD_low_ (Fig. 2c). Arg-rich sequences show higher transition midpoints for chain expansion than Lys-rich sequences, and Asp-rich sequences show slightly higher midpoints than Glu-rich sequences. Furthermore, at the highest salt concentrations probed — in the molar range — the Arg-rich sequences exhibit higher transfer efficiencies than the Lys-rich sequences. The stronger compaction of Arg-rich compared to Lys-rich variants at high salt highlights differences in the nature of the interactions of Lys and Arg, related to dissimilarities in their chemical structure, multivalency, charge delocalization, and polarizability, including a component of hydrophobicity induced at high salt^52-54^. The origin of the stronger intrachain interactions for Asp than for Glu is less pronounced, but possible contributions are differences in local interactions^47^, hydrophobicity^55^, or their interactions with salt ions^56^. Again, the resulting chain dimensions of the sequence variants at physiological salt concentrations are quite different (Fig. 2c). In summary, both the degree of charge segregation and the chemical identity of the charged residues impact the intramolecular interactions and chain dimensions of polyampholytic IDPs.

### Linking global observables to intramolecular interactions

To relate the experimental observables to conformational ensembles and intrachain interactions, we turn to molecular simulations, taking advantage of recent improvements^57^ in coarse-grained^12,31,58,59^ and atomistic^60-62^ force fields that provide increasingly accurate representations of IDPs. Our coarse-grained simulations use an HPS^31,58^ representation, where each residue is mapped to a bead with appropriate size and charge, and an interaction strength with other residues optimized with respect to experimental data^5,31,58,59,63^. Despite this simplicity, the resulting chain dimensions are often close to experiment^5,12,31^, and the moderate computational cost enables us to probe many sequences and salt concentrations. However, the absence of explicit solvent and the reduced number of degrees of freedom precludes a quantitative comparison of absolute timescales^64-66^. We thus complement coarse-grained with all-atom explicit-solvent simulations, which provide exquisite detail and absolute timescale information^60^, but where limitations in sampling lead to larger uncertainties, despite ~3 μs of simulation time for each variant and salt concentration. To enable a quantitative comparison with the FRET measurements, both models include explicit representations of the donor and acceptor dyes, previously optimized with respect to experiment^31,67,68^. The overall agreement between transfer efficiencies from experiment and simulations for our different sequence variants and salt concentrations results in concordance correlation coefficients of 0.95±0.01 for the coarse-grained and 0.81±0.07 for the atomistic simulations (Fig. 3a,b), respectively, illustrating the high quality of the models.

**Figure 3.**
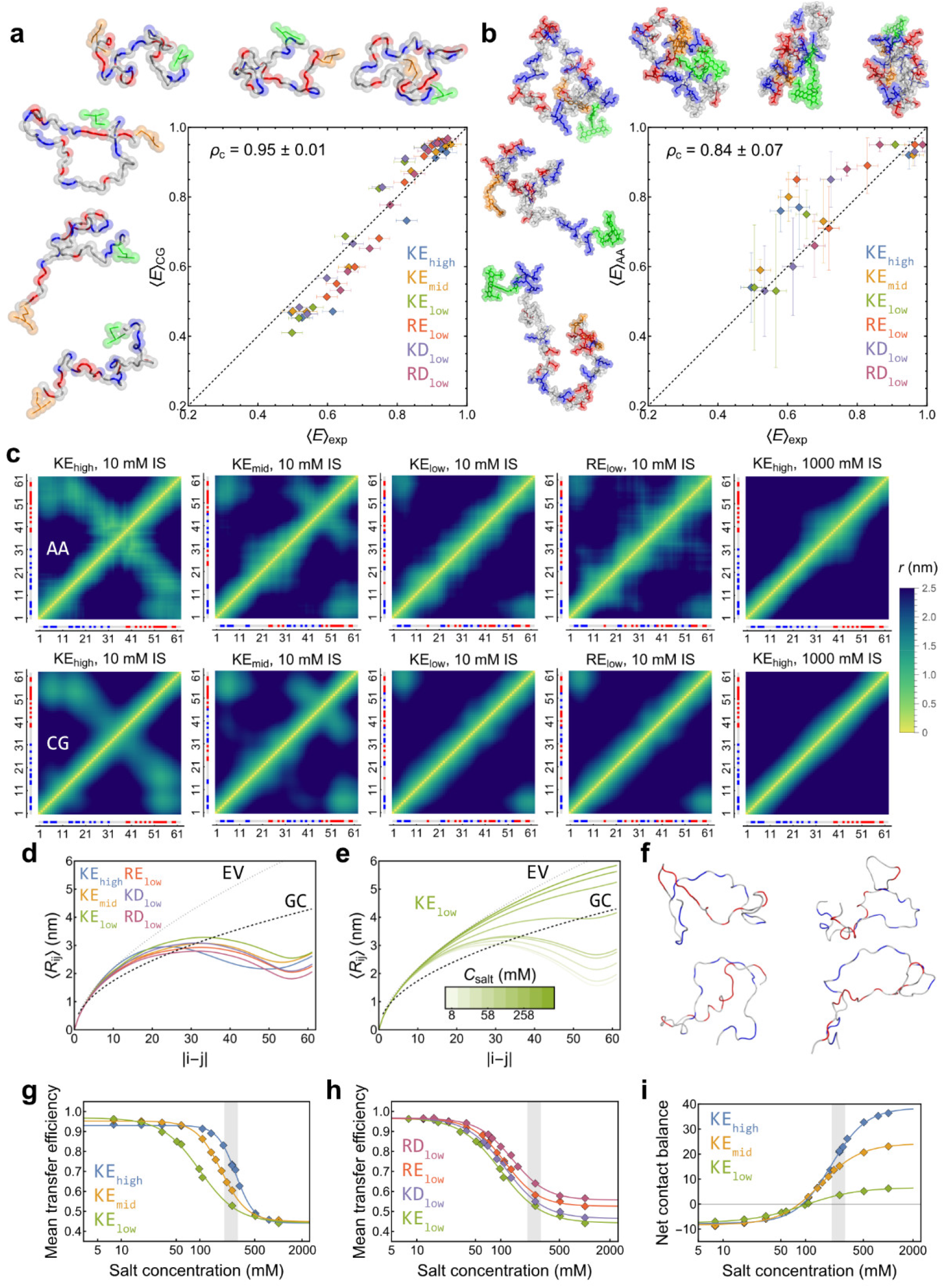
Simulations rationalize chain dimensions and intramolecular interactions. **a, b**, Comparison of ⟨E⟩ from experiment and coarse-grained MD (a) and all-atom MD simulations (b), with simulation snapshots illustrating the wide range of configurations sampled by KE_low_ (dashed line: identity line; error bars for MD: standard deviation of three independent runs or block averaging; error bars for experiment: systematic uncertainty due to instrument calibration; ρ_c_: concordance correlation coefficient). The slightly more compact configurations in all-atom MD than in experiment may originate from imperfections in the modeling of salt bridges.^71-73^ **c**, Inter-residue distance maps between Cα atoms for selected sequence variants at low and high salt concentration (ionic strength, IS) from all-atom (AA, top) and coarse-grained (CG, bottom) MD simulations. **d, e**, Average inter-residue distances, ⟨R_ij_⟩, as a function of sequence separation for different constructs (d) and salt concentrations (e) from coarse-grained simulations. The dependencies expected for Gaussian (dashed) and excluded-volume chains (dotted) are shown for comparison. **f**, All-atom MD snapshots illustrating the configurations sampled by KE_high_ (top) and KE_low_ (bottom) at 10 mM, highlighting the range of conformations sampled even at low salt concentration. **g-i**, Salt concentration dependence of mean transfer efficiency (g, h) and net contact balance (i) from coarse-grained simulations.

Intrachain distance maps (Fig. 3c) highlight the long-range attractive interactions within the more charge-segregated sequences that lead to their stronger compaction compared to less segregated sequences (Fig. 2c). The interaction patterns are remarkably similar for both simulation models. Electrostatic interactions are screened at high salt concentrations (Fig. 3c), and the resulting loss in long-range contacts causes the change in expansion that is observed experimentally (Fig. 2b,c). The balance between attractive non-local and repulsive local charge interactions, which are screened at different salt concentrations, influences the differences in transition midpoints we observe between the sequence variants. Especially the all-atom simulations also show slightly stronger attractive interactions for the Arg-rich compared to the Lys-rich sequences, predominantly for residues that are close in sequence (Fig. 3c). The influence of charge patterning on the conformational distributions is further reflected in the dependence of average inter-residue distances on sequence separation (Fig. 3d,e): In contrast to the simple monotonic dependence expected for homopolymers^69^, the patterned sequences exhibit the non-monotonic dependence characteristic of interactions that, on average, bring the chain ends close together^28,35^. Only at high salt concentrations, where charge interactions are screened and the chains expand, is a monotonic trend recovered. Notably, however, the conformational distributions obtained from the simulations show that even for the most charge-segregated variant, KE_high_, at the lowest salt concentrations, the relative arrangement of the long oppositely charged segments remains highly dynamic and disordered (Fig. 3f), in line with our CD and NMR results (Fig. 1), and far from a persistent “ladder-like” arrangement sometimes envisioned for very strong polyelectrolyte interactions^70^. In summary, the simulations allow us to rationalize the strong effects of sequence composition, charge segregation, and salt concentration on the compaction and conformational distributions of polyampholytic IDPs in terms of intrachain interaction patterns. But how are the observed transitions related to the underlying free-energy surfaces?

### Cooperativity and free-energy surfaces of charge-patterned IDPs

The coarse-grained simulations reproduce not only the experimentally observed chain expansion and the shift in transition midpoints with increasing charge segregation (Fig. 2b,c), but also the concomitant increase in the steepness of the transitions (Fig. 3g-h). The sigmoidal transitions and the increasing steepness with charge segregation can be considered evidence for cooperativity^74^. However, in contrast to the two-state cooperativity observed in processes such as protein folding^75,76^, both the single-molecule FRET experiments (Fig. 2a) and the simulations suggest a continuous transition of the conformational ensembles shifting from compact to expanded (Fig. 4c), rather than an interconversion between two well-defined thermodynamic states. Even so, since the amino acids are linked in the polypeptide chain, their interactions are not independent, because they are coupled with those of neighboring residues and thus cooperative in the sense of this collective behavior: If one pair of oppositely charged residues make an interaction, the neighboring residues are also brought into the local contact environment by backbone connectivity. Accordingly, the more charge-segregated a sequence is, the more attractive interactions form simultaneously^77^ (Fig. 4d), leading to stronger chain compaction. Even if no secondary or tertiary structure is formed (Fig. 1e-h), the salt-dependent transitions of charge-segregated IDPs can thus be cooperative, as previously suggested based on simulations^21,78,79^.

**Figure 4.**
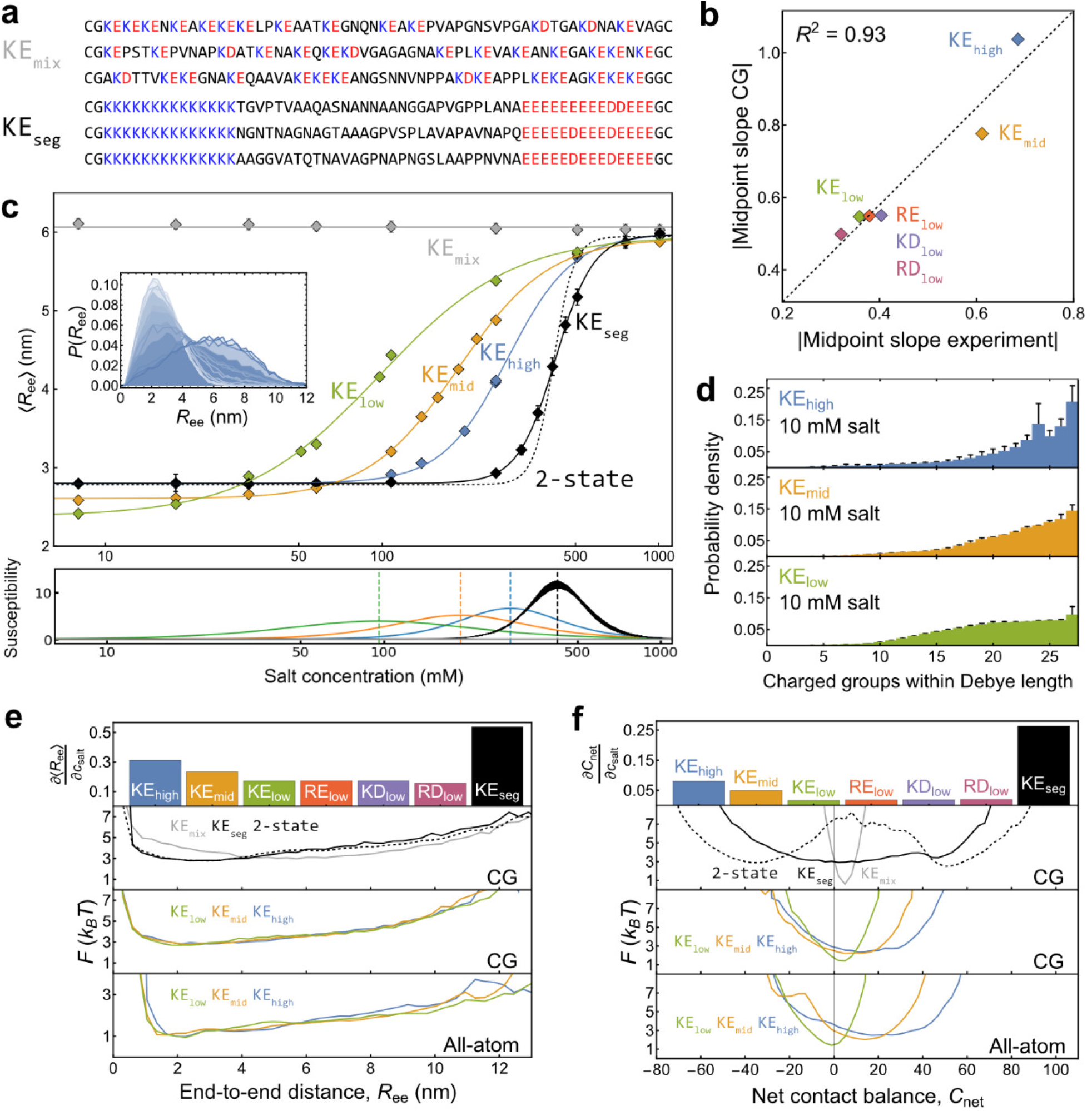
Quantifying the relation between charge segregation and cooperativity from experimentally validated simulations. **a**, Examples of sequences for KE_mix_ (upper) and KE_seg_ (lower), with negatively charged residues highlighted in red and positively charged ones in blue. **b**, Comparison between slopes at the midpoints for transitions based on the mean transfer efficiency as a function of salt concentration, a⟨E⟩/aIogc_saIt_, for the different sequence variants (cf. Figs. 2b,c and 3g,h; see legend for color code; if error bars are not visible, they are smaller than the symbols). **c**, (top) Average dye-to-dye distance, ⟨R_ee_⟩, from coarse-grained simulations as a function of salt concentration for the different sequence variants (see legend for color code, inset: end-to-end distance as a function of salt concentration). (bottom) Corresponding susceptibilities calculated from the slopes of the fits to the transitions (see Methods). **d**, Contact probability distributions for different sequence variants show that greater charge segregation leads to a larger number of charge contacts per residue. **e**, (top) Midpoint slopes for transitions based on ⟨R_ee_⟩ as a function of salt concentration for the different sequence variants. Free-energy profiles for R_ee_ as a reaction coordinate at the transition midpoints from coarse-grained (CG) and all-atom simulations (All-atom). See legend for sequence variants. **f**, Analogous to (e) for the net contact balance, C_net_, as a reaction coordinate.

How can we quantify the degree of cooperativity and relate it to the shape of the free energy surface in such systems? Models commonly used to quantify cooperativity in the absence of simple two-state behavior, such as Hill^80^ and Zimm-Bragg^81^ theory are difficult to apply, since in our case there are no well-defined binding sites or individual structurally defined interactions. However, a quantity that is related to cooperativity in all these scenarios is the variance of an observable or order parameter, a measure of the equilibrium fluctuations in a system. A familiar example is the heat capacity, which corresponds to the magnitude of energy fluctuations that determine the thermal response, with a maximum near phase transitions or conformational changes^82-84^. In cooperative ligand binding, the Hill slope can be expressed as the variance of the number of bound ligands normalized by the binomial variance expected for independent binding sites^85,86^. Even for a two-state process, the variance of an observable whose values differ between the two states is maximal at the transition midpoint^84^. For suitably chosen order parameters, the variance can be related to a generalized susceptibility^82,87^ that quantifies the magnitude of the response of the order parameter to a perturbation. In our case, the susceptibility would correspond to the change in chain dimensions or intrachain contacts in response to an increase in salt concentration, given by the slope of the transition curve, which is maximal at the midpoint (Fig. 4c). Owing to its relation to the variance, the susceptibility reflects the magnitude of fluctuations in intrachain interactions or chain dimensions^88^ (see Methods for details). In summary, the variance and susceptibility thus provide generalized measures of cooperativity that include two-state and Hill-type behavior as special cases. We will use them here to assess the influence of charge patterning on the cooperativity of salt-induced transitions of disordered polypeptides.

Both in experiment (Fig. 2b) and in the coarse-grained simulations (Fig. 3g), the slope at the transition midpoint for the transfer efficiency as a function of salt concentration increases with increasing charge segregation. Even quantitatively, the values are highly correlated and very similar (Fig. 4b). In contrast to the experimental data, which only yield information on a small number of collective order parameters, such as the average end-to-end distance, the experimentally validated simulations provide access to the complete set of chain configurations, or microstates, which enables a much more comprehensive analysis in terms of statistical mechanics. To be able to put the cooperativity values based on susceptibility into context, we first identify two sets of reference sequences that represent limiting values of cooperativity. The lowest cooperativity is represented by sequences with the same amino acid composition as KE_low_, KE_mid_, and KE_high_, but without long-range charge interactions, which we achieve by distributing KE pairs uniformly along the chain (KE_mix_, Fig. 4a). The coarse-grained simulations of 128 representative sequences exhibit virtually no change in the conformational ensemble with salt concentration, corresponding to a cooperativity close to zero (Fig. 4c). Among the sequences with the same composition, the highest cooperativity is achieved by clustering the positive and negative charges at opposite chain termini (KE_seg_, Fig. 4a), which indeed results in much steeper transitions (Fig. 4c). Finally, the upper limit on the cooperativity scale can be represented by a hypothetical two-state-like system composed of two fixed conformational ensembles, the ones at 8 mM and 1 M ionic strength of the KE_seg_ sequences; all average values of the desired observables, such as the average end-to-end distance, are then computed based on a linear combination of these two states, conceptually similar to the approach used for two-state protein folding^75,76^, with the relative populations determined by their screened Coulomb energies (Fig. 4c, see Methods for details). While the underlying assumption that the two individual ensembles do not change with salt concentration is not realistic, this scenario corresponds to the steepest transition if we assume that the ionic strength-dependence arises only from the shift in populations due to the change in electrostatic energy of the system. If we define the cooperativities of the two limiting references as 0 (KE_mix_) and 1 (hypothetical two-state), respectively, we can use the midpoint slopes observed for our variants with different charge segregation for quantifying their cooperativity on this scale (Fig. 4e,f).

We illustrate this analysis for two representative order parameters: On the one hand, the end-to-end distance, *R*_ee_, a parameter closely related to the experimentally observed transfer efficiency (Fig. 4e); and on the other hand, the net electrostatic contact balance, *C*_net_ = *C*_like_ – *C*_±_, the difference between the number of contacts between like-charged, *C*_like_, and oppositely charged residues, *C*_±_, a parameter that is mechanistically more informative regarding the interesting interplay of attractive and repulsive interactions within the chain^28^, but that is only available from the simulations (Fig. 4f). The resulting absolute cooperativity values differ between the two order parameters, but the relative values show the same trends, with a decrease in cooperativity in the order KE_high_ > KE_mid_ > KE_low_, and similar cooperativity for KE_low_, RE_low_, KD_low_, and RD_low_.

Based on the simulations, we can relate the cooperativity of the different sequences to their free-energy surfaces. We focus on the transition midpoints, where differences between sequence variants are expected to be most pronounced. Again, we illustrate the results with *R*_ee_ and *C*_net_ as order parameters for KE_high_, KE_mid_, and KE_low_, the two reference sequences, KE_mix_ and KE_seg_, and the hypothetical two-state case (Fig. 4e,f). For *R*_ee_, the free-energy surfaces at the midpoint are broad and barrierless in all cases, and the differences in width for the different sequences are too small to be visually noticeable, since they are dominated by the broad distance distributions of the expanded configurations of the chains (Fig. 4e, Fig. S4e). We thus focus on *C*_net_, for which the differences are much more pronounced. The limiting hypothetical two-state scenario yields a barrier of ~4 *k*_B_*T*. However, even for KE_seg_, we obtain a potential with a very small barrier of less than 1 *k*_B_*T*. For the experimentally investigated sequences KE_low_, KE_mid_, and KE_high_, no barriers are detectable, in line with the single, continuously shifting FRET efficiency populations and NMR resonances observed experimentally (Figs. 1f, 2a). This simulation result is not caused by the simplicity of the coarse-grained model or the choice of order parameter; atomistic simulations also show no barriers (Fig. S5), even with optimized reaction coordinates based on principal component or autoencoder analysis (Fig. S6, S7). Residual ‘roughness’ in the free-energy surfaces is below 1 *k*_B_*T* and at least in part originates from the limited sampling in the all-atom simulations.

How can barrierless free-energy surfaces be compatible with cooperativity? The answer goes back to our discussion of the variance and is rooted in relations that link the response upon perturbation to the magnitude of equilibrium fluctuations^82,89^. For instance, the free-energy surface along *C*_net_ is much wider for KE_high_ than for KE_low_ (Fig. 4f), indicating larger variance, and thus greater equilibrium fluctuations. As a result, a smaller perturbation in terms of a change in salt concentration is sufficient to shift the ensemble for KE_high_ than for KE_low_, corresponding to a larger slope of the observable as a function of salt concentration at the midpoint, i.e., a greater susceptibility. Even flat and barrierless free-energy surfaces can thus encode cooperativity. It is worth noting that the widths of the free-energy surfaces along *R*_ee_ and *C*_net_ — and thus the variances — exhibit very different sensitivity to charge segregation (Fig. 4e,f, Fig. S4). The reason is that salt acts primarily by screening electrostatic interactions, so the most sensitive reaction coordinates are ones like *C*_net_ that track the electrostatic contacts that form or dissolve during expansion; such coordinates thus respond directly to the molecular changes that distinguish highly charge-segregated sequences from more mixed sequences. *R*_ee_, on the other hand, is dominated by large fluctuations unrelated to salt-dependent rearrangements. The reason why ⟨*R*_ee_⟩ as a function of salt concentration still shows a pronounced transition with clear differences between sequence variants (Fig. 4c) is that the susceptibility to ionic strength does not depend on the variance of *R*_ee_; instead, it results from the covariance with a generalized force conjugate to the control parameter, a quantity related to the energy of the configurations (see Methods for details; Fig. S8). More intuitively, the values of ⟨*R*_ee_⟩ at low and high salt, respectively, are very similar for the different sequence variants, so the variances of ⟨*R*_ee_⟩ at the respective midpoints are also similar. However, the susceptibility further depends on how sensitively the weights of the different configurations respond to salt concentration, i.e., on the energetics of their electrostatic contact patterns.

In summary, charge-patterned IDPs can exhibit substantial differences in cooperativity in salt concentration-dependent transitions, even though no secondary or tertiary structure is formed or broken. The final question we want to address is how their equilibrium behavior and the flat free energy surfaces relate to the dynamics of such IDPs.

### Effect of charge patterning on chain dynamics

The presence of single FRET-efficiency peaks and NMR resonances shifting continuously as a function of salt concentration (Figs. 1f, 2a) indicate fast exchange among all configurations relative to the millisecond observation times of these experiments^44^. Burst variance analysis^90^ confirms the absence of FRET efficiency peak broadening beyond shot noise (Fig. 5a). These observations are consistent with the lack of secondary or tertiary structure formation (Fig. 1e-h) and with the absence of free-energy barriers between compact and expanded configurations observed in the simulations (Fig. 4e,f, Fig. S5, S6, S7). To directly measure the time scale on which our polyampholytic IDPs explore their free energy surfaces, we use nanosecond fluorescence correlation spectroscopy (nsFCS)^91^. This approach allows us to monitor the distance fluctuations between the donor and acceptor dyes attached to the chain ends down to the low nanosecond range and thus provides a sensitive way of probing the conformational dynamics of IDPs^91-96^.

**Figure 5.**
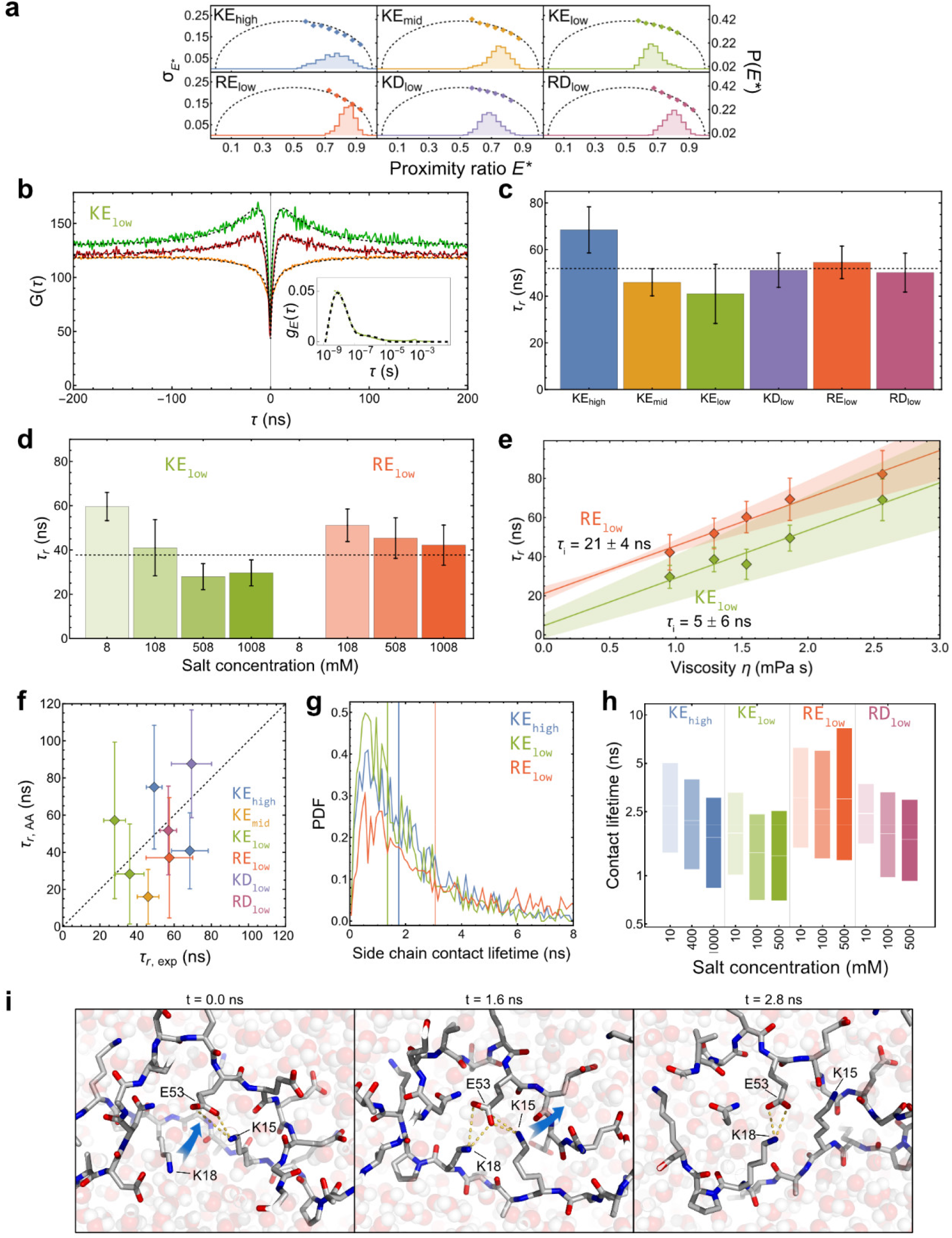
Rapid chain dynamics of charge-patterned IDPs. **a**, Burst variance analysis^90^ at the midpoint of each protein variant (colored lines: experimental proximity ratio (E*) histograms; dashed lines: burst variance expected from shot noise; colored diamonds: observed burst variance). **b**, Nanosecond fluorescence cross-(orange) and autocorrelations of the donor (green) and acceptor (red) for KE_low_ at 100 mM salt (fit: dashed lines, see Methods for details). Inset: FRET efficiency correlation^97^ of the same data with fit (see Methods for details). **c**, Reconfiguration times, τ_r_, from the FRET efficiency correlations of all sequence variants at their respective midpoints. **d**, τ_r_ of KE_low_ and RE_low_ from FRET efficiency correlations at different salt concentrations. **e**, Assessing internal friction from solvent viscosity-dependent τ_r_ of KE_low_ and RE_low_ from FRET efficiency correlations at 1000 mM salt concentration (lines: linear fits; error bars from bootstrapping, see Methods). **f**, Comparison of τ_r_ from experiment and all-atom MD simulations at the midpoint for the different sequence variants (see (c) for color code; dashed line: identity line; error bars for MD: standard deviation from three independent runs; error bars for experiment, see Methods). **g**, Probability density functions of contact lifetimes for KE_high_, KE_low_, and RE_low_ at 10 mM KCl from all-atom simulations (see Methods) (vertical lines: median). **h**, Box chart of contact lifetimes from all-atom simulations for several variants at different salt concentrations (box boundaries: 25th and 75th percentiles; white line: median). **i**, Illustration of dynamic shuffling between side chain contacts via ternary interactions from all-atom MD.

We observe end-to-end distance dynamics in the tens-of-nanoseconds range, as indicated by the simultaneous relaxation components in the cross-and autocorrelations with opposite sign, the characteristic signature of distance dynamics^24,91^ (Fig. 5b). From a global analysis of all three correlation functions, we obtain the end-to-end distance correlation time, or reconfiguration time (see Methods for details). We also used the recently developed analysis in terms of FRET efficiency correlations^97^, which yields similar results (Fig. 5b). For all variants and salt concentrations, we observe reconfiguration times, *τ*_r_, between 30 ns and 70 ns (Fig. 5c,d), typical of IDPs of this length^24^. Owing to the presence of microsecond triplet blinking in the Alexa fluorophores^98,99^, it is difficult to exclude the presence of dynamics in this time range from these measurements alone. We thus employed a second dye pair with lower triplet populations, Cy3B-CF660R^100^, and detect no anticorrelated component in the microsecond range that would indicate distance dynamics on this timescale (Fig. S9). Together with the continuous shifts of the FRET efficiency populations as a function of salt, indicating fast exchange between conformations on the burst time of ~0.1 ms (Figs. 2a), we thus have no experimental evidence for conformational dynamics slower than the tens-of-nanosecond timescale corresponding to elementary chain reconfiguration, even close to the midpoints. Together with the observation that the chain reconfiguration times are close to expectations for disordered proteins of this length lacking charge patterning^24^, these results are consistent with the absence of free-energy barriers observed in the simulations (Fig. 4e,f).

It may be surprising that the chain dynamics are so rapid, even for sequences with large charge segregation, and even at the lowest salt concentrations (Fig. 5d), where the strength of the Coulomb forces might be expected to cause a pronounced increase in intrachain interactions that could lead to a substantial slow-down of chain reconfiguration. To elucidate the origin of the observed rapid chain dynamics, we turn to the all-atom explicit-solvent MD simulations, since they provide absolute timescale information. Although extensive sampling of conformational space is challenging in these computationally expensive simulations, the resulting dye-to-dye distance relaxation times are in the correct range compared to experiment (Fig. 5f). Together with the similarity of transfer efficiencies between experiment and simulation (Fig. 3b), we conclude that the Amber99SBws^60^ force field provides a reasonable representation of the dynamics in our polyampholytic IDPs. The MD simulations suggest a solution to the conundrum of rapid dynamics: The lifetimes of side chain contacts within the chain are remarkably short, in the low nanosecond range (Fig. 5g), and exhibit only a moderate increase with decreasing salt concentration (Fig. 5h), in line with the concomitant increase in *τ*_r_ for KE_low_ and RE_low_ (Fig. 5d). Besides the high level of solvation of the charged side chains^101,102^, an important contribution to these short contact lifetimes is the long-range nature of electrostatic interactions, which enables a rapid exchange between contact pairs via transient ternary interactions between side chains: For the example illustrated in Fig. 5i, starting from an initial contact between a Lys and a Glu sidechain, another Lys sidechain makes a contact and replaces the original Lys. Indeed, in the context of such highly charged sequences in their compact ensembles at low salt concentration, not only a few but many charged residues are within a Debye length of each charged group (Fig. 4d). The interaction among all these residues may be more adequately described by a mean field-type interaction^103^ than by well-defined individual salt bridges. Similar behavior has previously been reported in phase-separated biomolecular condensates between H1 and ProTα^100,104^ and may be a more general mechanism that prevents kinetic arrest in assemblies of highly charged biomolecules.

Based on the contact lifetimes, we noticed that the simulations suggest a slight difference in dynamics between Lys-and Arg-rich sequences: For RE_low_, the contact lifetimes are consistently higher than for KE_low_, and the difference persists even at the highest salt concentrations (Fig. 5h), pointing towards a residue-specific effect on chain dynamics, in line with the greater compaction of the Arg variants at high salt concentrations (Fig. 2c). Prompted by these results, we performed nsFCS measurements to assess the possible role of internal friction^92,105-107^ in RE_low_ at 1 M KCl. The commonly employed experimental strategy to quantify internal friction in IDPs is to measure the chain reconfiguration time, *τ*_r_, as a function of solvent viscosity. The solvent-independent contribution to friction, the internal friction time, *τ*_i_, is then obtained by extrapolating to zero solvent viscosity^24,92^ (Fig. 5e). We indeed observe a difference between RE_low_ and KE_low_: The values of *τ*_r_ are slightly but systematically greater for RE_low_ than for KE_low_ at all viscosities. For KE_low_, the resulting value of *τ*_i_ is 5 ± 6 ns, i.e., indistinguishable from zero within experimental uncertainty, indicating the absence of internal friction. For RE_low_, however, *τ*_i_ = 21 ± 4 ns, suggesting that the slightly slower chain dynamics observed at high salt concentrations both in simulation and experiment are caused by internal friction, possibly owing to a contribution from hydrophobic interactions of Arg residues. At low salt concentrations, the difference in *τ*_i_ between RE_low_ and KE_low_ is still present but smaller (Fig. S10).

## Discussion

In summary, we observe pronounced effects of charge patterning on the conformational distributions of disordered polyampholytic proteins and on their salt concentration dependencies. Increasing charge segregation leads to stronger long-range attraction between the oppositely charged regions of the chain, as reflected by a larger resistance to salt-induced expansion and more cooperative transitions. Notably, these differences in cooperativity occur despite the absence of free-energy barriers. To quantify cooperativity, we thus employ a generalized concept based on the susceptibility, which peaks at the transition midpoint and relates the fluctuations in a system to its response to a perturbation, such as the salt concentration: More cooperative sequence variants exhibit larger equilibrium fluctuations at the midpoint, visible in simulations as broader free-energy surfaces along suitable reaction coordinates; as a result, less of the perturbation is required to shift the conformational ensemble from compact to expanded, corresponding to steeper transitions. In keeping with the flat free-energy surfaces, the chain dynamics are very rapid for all sequence variants, in the range between 30 and 70 ns, as expected for IDPs without charge patterning.^24,92,108^ The chain dimensions and the response to salt concentration are also influenced by the identity of the charged residues. Especially Arg mediates stronger intrachain interactions than Lys and results in more compact configurations even at molar salt concentrations, highlighting the differences in the nature of the interactions of the two basic side chains^52,109-111^. Similarly, Arg facilitates the formation of biomolecular condensates by phase separation^112-114^, including complex coacervation between ProTα and Arg-rich IDPs^104^. The difference between the acidic side chains is less pronounced but still detectable, with slightly stronger interactions for Asp than for Glu.

The cooperativity we observe for polyampholytic IDPs with SCD values in the naturally occurring range is much lower than for pronounced two-state systems with a large barrier, such as the barrier between folded and unfolded states in many proteins that form a well-defined folded structure. For such two-state folders, the slope of the folding transition (‘m-value’) is correlated with the decrease in solvent-accessible surface area upon folding and thus with the number of interactions formed in the folded structure^115^. The disordered proteins we investigate here remain highly solvated even in their most compact configurations, so the concept of surface accessible surface area is difficult to apply. However, they behave similarly in the sense that the variants that show higher cooperativity tend to make a large number of intrachain interactions at low salt concentration, i.e., a larger number of charged residues are located within one Debye length (Fig. 4d), corresponding to a larger cluster of cooperatively interacting charges. Another interesting link is that cooperativity in protein folding is related to the change in heat capacity between folded and unfolded states^115^, which in turn is related to the magnitude of the energy fluctuations near the unfolding midpoint^82^. Analogously, the cooperativity we observe as a function of salt concentration can be related to the fluctuations in free energy near the midpoint, as reflected in the variance of the conformational ensemble and the width of the underlying free energy surfaces (Fig. 4f, Fig. S8).

Our results illustrate the growing synergy between experiment and simulation, where experimental data serve as benchmarks for the simulations, and the simulations enable a detailed interpretation of measurements at the molecular level. This approach benefits from the complementarity of coarse-grained and all-atom simulations as well as analytical theory^116^ to extend our quantitative understanding of IDPs beyond their conformational distributions to their dynamic properties. In the present case, on the one hand, coarse-grained simulations, which allow for a more extensive sampling of the distance distribution, enable us to probe a large number of solution conditions and sequence variants and address the question of cooperativity in polyampholytic IDPs quantitatively (Fig. 4). On the other hand, while all-atom explicit-solvent simulations are more difficult to converge, they can provide absolute timescale information (Fig. 5). Since the chain reconfiguration times from the simulations are close to the experimental values, the dynamics of interactions within the chain are likely to be represented reasonably accurately.

For our charged sequence variants, the contact lifetimes observed in the all-atom simulations remain in the low nanosecond range under all conditions, even at the lowest salt concentrations. These exceedingly short-lived interactions are likely to be caused by a high level of side chain hydration^101,102^, but also by the rapid exchange of electrostatic interactions between oppositely charged residues via ternary or higher-order contacts in the large clusters of charged residues within their compact conformational ensembles, similar to the behavior observed in complex coacervates of charged IDPs^100,104^. A likely functional consequence of these fast chain dynamics is that the rapid downhill binding kinetics observed for highly charged IDPs^6,44^ can occur irrespective of charge patterning. Evolutionary variation in charge patterning thus enables pronounced changes in intramolecular distance distributions without compromising the rapid dynamics of these systems, which may be an important factor for polyampholytic sequences, which are highly prevalent in disordered proteins^12^.

## Methods

### Protein expression

^15^N-labeled ProTα was produced as previously described^5^. Codon-optimized protein sequences including terminal cysteines for site-specific labeling were obtained from GeneCust (Boynes, France). Protein sequences were cloned into pET-20b(+) plasmids (EMD Millipore) with a C-terminal SUMO domain for enhanced expression. This domain was fused to a His_6_-tag, which was separated from the linker sequence of interest via an Ulp1 cleavage site. All protein constructs were expressed in Rosetta DE3 (EMD Millipore). Cultures were grown to an OD_600_ of 1 in 2xYT medium (Sigma-Aldrich) containing kanamycin, induced with 1 mM isopropyl β-D-1-thiogalactopyranoside and then incubated at 20°C for a period of 12 hours. The cells were harvested, the resulting pellets resuspended in lysis buffer (100 mM NaH_2_PO_4_/Na_2_HPO_4_, 10 mM Tris-HCl, 6 M guanidinium chloride (GdmCl), 10 mM imidazole, 1 mM dithiothreitol (DTT), pH 8.0) and left to incubate at 4°C overnight with shaking. The soluble fraction was purified by nickel chelate affinity chromatography (Ni Sepharose Excel Cytiva), where lysis buffer was used for washing. The elution buffer composition was as follows: 100 mM NaH_2_PO_4_/Na_2_HPO_4_, 10 mM Tris-HCl, 6 M GdmCl, 500 mM imidazole, pH 8.0. The samples were then dialyzed against 50 mM Tris-HCl, 150 mM NaCl, 10% (v/v) glycerol, and pH 8.0. Subsequently, Ulp1 was added at a concentration of 10 U/mg, and the proteolytic reaction was allowed to proceed for 3 hours at room temperature. The reaction was quenched by the addition of 1 g/ml GdmCl. The protein solutions were concentrated to a total volume of approximately 1 mL using an Amicon Ultra Centrifugal Filter, 3 kDa MWCO (EMD Millipore).

For the study of ^13^C-^15^N labeled proteins, an M9 minimal medium was utilized, comprising 45 mM NaHPO_4_, 22 mM KH_2_PO4, 8.5 mM NaCl, 18.5 mM ^15^NH_4_Cl, 3 g/L ^13^C glucose, 2 mM MgSO_4_, 10 μM CaCl_2_, 1 mg/L biotin, 5 mg/L thiamine, 50 μg/mL kanamycin, and 10 mL/L of trace-element solution. The solution of trace elements was composed of the following substances, all dissolved in deionized water: 20 μM ethylenediaminetetraacetic acid, 3 mM FeCl3, 600 μM ZnCl^2^, 76 μM CuCl^2^, 77 μM CoCl^2^, 161 μM H3BO3, and 43 μM MnCl^2^. The pH of the solution was adjusted to 7.5. A 20 mL preculture was grown in LB medium (Sigma-Aldrich), with the addition of kanamycin. Upon reaching an OD_600_ of 1, 1 mL of the preculture was utilized to inoculate 100 mL of preculture with M9 medium. Subsequently, 50 mL of the last preculture were utilized to inoculate 1 L of M9 medium, and 1 mM isopropyl β-D-1-thiogalactopyranoside was added upon reaching an OD_600_ of 1. The subsequent steps in this procedure were analogous to those previously outlined for the non-isotope-labeled samples.

### Protein Labeling

Protein samples were reduced by adding DTT to a final concentration of 10 mM. The purification of the constructs was subsequently conducted via reversed-phase high performance liquid chromatography (RP-HPLC) on a C18 column (Dr. Maisch Reprosil Gold 200 C18, 5 µm) with 0.1% (v/v) trifluoroacetic acid (aq) as buffer A and acetonitrile as buffer B. The eluates were then lyophilized overnight and subsequently re-dissolved in 50 mM NaH_2_PO_4_/Na_2_HPO_4_, 6 M GdmCl, pH 7.4. Protein concentrations were quantified based on their absorbance at 205 nm (extinction coefficient 31 mL mg^-1^ cm^-1^). For the fluorophore labeling procedure, Alexa Fluor 488 C5 maleimide (Thermo Fisher Scientific) or Cy3B maleimide (GE Healthcare) was dissolved in anhydrous dimethylformamide, and the solution was added at a molar ratio of protein to dye of 1:0.7. The reaction was permitted to continue at a temperature of 4 °C overnight, after which it was quenched by the addition of DTT at a final concentration of 10 mM. Singly labeled protein was separated from unreacted and doubly labeled protein by RP-HPLC (see above), followed by lyophilization. Donor-labeled protein was re-dissolved in a buffer composed of 50 mM NaH_2_PO_4_/Na_2_HPO_4_, 6 M GdmCl, and pH 7.4. Alexa Fluor 594 C5 maleimide (Thermo Fisher Scientific) or CF660R maleimide (Biotium) was dissolved in anhydrous DMSO, and the acceptor dye added to the donor-labeled protein solution at 3-fold molar excess. The reaction was permitted to proceed at 4 °C overnight and then quenched by adding DTT to a final concentration of 10 mM. The purification of the donor-acceptor labeled protein was achieved through RP-HPLC (see above), followed by lyophilization and re-dissolution in 50 mM NaH_2_PO4/Na_2_HPO4, 6 M GdmCl, pH 7.4. The protein identity and site-specific labeling were confirmed by electrospray mass spectrometry. Fluorescently labeled protein constructs were stored at -80 °C until further use.

### Single-molecule spectroscopy

Single-molecule fluorescence experiments were performed on a four-channel MicroTime 200 confocal instrument (PicoQuant) equipped with an Olympus UplanApo 60x/1.20 Water objective. Alexa 488 was excited with a diode laser (LDH-D-C-485, PicoQuant) at an average power of 100 µW (measured at the back aperture of the objective). The laser was operated in continuous-wave mode for nsFCS experiments and in pulsed mode with interleaved excitation^117^ (PIE) for fluorescence lifetime measurements. The wavelength range used for acceptor excitation was selected with two band pass filters (z582/15 and z580/23, Chroma) from the emission of a supercontinuum laser (EXW-12 SuperK Extreme, NKT Photonics) operating at a pulse repetition rate of 20 MHz (45 µW average laser power after the band pass filters). The SYNC output of the SuperK Extreme was used to trigger interleaved pulses from the 488-nm diode laser. Sample fluorescence was collected by the microscope objective, separated from scattered light with a triple band pass filter (r405/488/594, Chroma) and focused on a 100-µm pinhole. After the pinhole, fluorescence emission was separated into two channels, either with a polarizing beam splitter for fluorescence lifetime measurements, or with a 50/50 beam splitter for nsFCS measurements to avoid the effects of detector deadtimes and afterpulsing on the correlation functions. Finally, the fluorescence photons were distributed by wavelength into four channels by dichroic mirrors (585DCXR, Chroma), additionally filtered by band pass filters (ET 525/50 M and HQ 650/100, Chroma), and focused onto one of four single-photon avalanche detectors (SPCM-AQRH-14-TR, Excelitas). The arrival times of the detected photons were recorded with a HydraHarp 400 counting module (PicoQuant, Berlin, Germany). All single-molecule experiments were conducted on freely diffusing molecules in 18-well plastic slides (ibidi) at 22 °C with concentrations of labeled protein between 50 and 250 pM in 10 mM Tris buffer pH 7.4, 0.01% Tween 20, 143 mM β-mercaptoethanol, with different concentrations of KCl for the salt titrations, and with appropriately chosen concentrations of glycerol for the viscosity dependence.

### Single-molecule FRET data analysis

Data analysis was carried out using the Mathematica (Wolfram Research) package Fretica (https://github.com/SchulerLab). For the identification of photon bursts, the photon recordings were time-binned at 1 ms. Photon numbers per bin were corrected for background, crosstalk, differences in detection efficiencies, and quantum yields of the fluorophores, and for direct excitation of the acceptor^118^. Bins with more than 30 photons were identified as photon bursts. Ratiometric transfer efficiencies were obtained for each burst from:

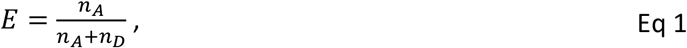

where *n*_A_ and *n*_D_ are the corrected numbers of donor and acceptor photons in the photon burst, respectively. The *E* values were histogrammed. The subpopulation corresponding to the FRET-labeled species was fitted with a Log-normal peak function to obtain the mean transfer efficiency, ⟨*E*⟩. Bursts from experiments in PIE mode were further selected according to the fluorescence stoichiometry ratio:

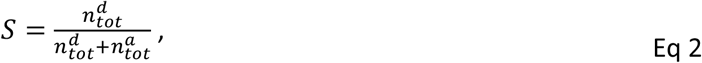

where 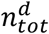 is the total number of photons after donor excitation and 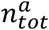 after acceptor excitation. The following relation was used to infer the end-to-end distance distribution, *P*(*r*) from the experimentally determined mean transfer efficiency, ⟨*E*⟩:^24^

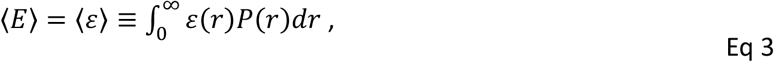

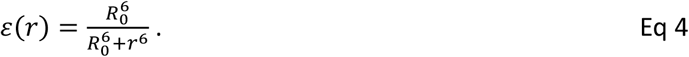

To this end, both the distance distribution of a self-avoiding walk polymer and the distance distribution obtained by all-atom molecular dynamics simulations were utilized for *P*(*r*). For the self-avoiding walk polymer, we use the SAW-v distance distribution^119^:

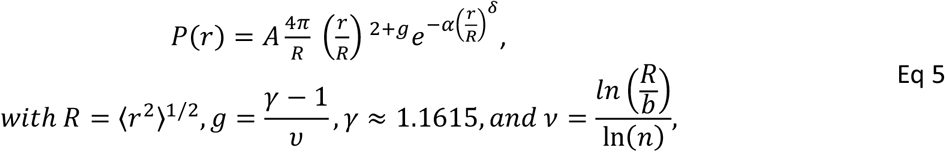

where *b* and *n* are the segment length and the number of segments of the polymer, respectively. The constants *A* and *α* are determined, for given values of *v* and *R*, from the conditions 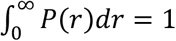 and 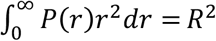, and *δ* = 1/(1 − *v*). In the context of our system, the value of *n* was set to 66, thereby accounting for the dye linkers^120^. *b* was set to 0.55 nm, as previously estimated for unfolded proteins^27,119^.

### Fluorescence correlation spectroscopy (FCS)

For FCS, the same experimental conditions as described in “Single-molecule spectroscopy” were used. The correlation between two time-dependent signal intensities, *I*_*i*_(*t*) and *I*_*j*_(*t*), measured on two detectors *i* and *j*, is defined as:

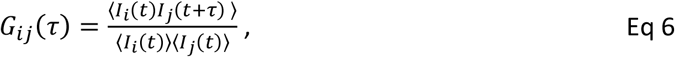

where the pointed brackets indicate averaging over *t*. In our experiments, we use two acceptor and two donor detection channels, resulting in the autocorrelations *G*_*AA*_(*τ*) and *G*_*DD*_(*τ*), and cross-correlations *G*_*AD*_(*τ*) and *G*_*DA*_(*τ*). By correlating detector pairs, and not the signal from a detector with itself, contributions to the correlations from dead times and afterpulsing of the detectors are eliminated^91,121^. Full FCS curves with logarithmically spaced lag times ranging from nanoseconds to seconds were fitted with^122,123^

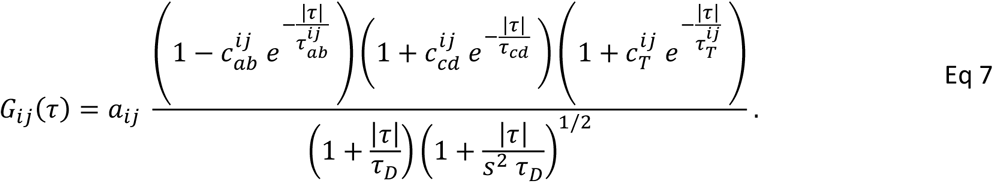

The three terms in the numerator with amplitudes *c*_*ab*_, *c*_*cd*_, *c*_*T*_, and timescales *τ*_*ab*_, *τ*_*cd*_, *τ*_*T*_ describe photon antibunching, chain dynamics, and triplet blinking, respectively. *τ*_*D*_ is the translational diffusion time of the labeled molecules through the confocal volume; a point spread function (PSF) of 3-dimensional Gaussian shape is assumed, with a ratio of axial over lateral radii of *s* = *ω*_z_/*ω*_xy_ (*s* = 5.3), and *a*_*ij*_ is the amplitude of the correlation functions. Parameters without indices *ij* are treated as shared parameters in the global fits of the auto- and cross correlation functions. To study the dynamics in more detail, donor and acceptor fluorescence auto- and cross correlation curves were computed and analyzed over a linearly spaced range of lag times, *τ*, up to a maximum, *τ*_max_, that exceeds *τ*_*cd*_ by an order of magnitude. For the subpopulation-specific analysis, we used only photons of bursts with *E* in the range of ±0.2 of the mean transfer efficiency of the FRET-active population, which reduces the contribution of donor-only and acceptor-only signal to the correlation. For direct comparison, correlation curves were normalized to unity at *τ*_*max*_. After normalization and in the limit of 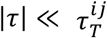 and 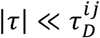, eq. 7 reduces to

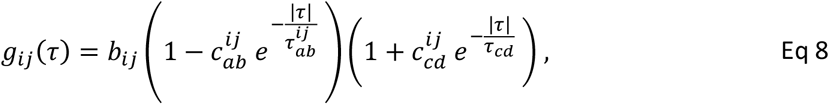

where *b*_*ij*_ = 1/*G*_*ij*_(*τ*_max_) is a normalization constant.

FRET efficiency correlation analysis was performed as described by Terterov et al.^97^ The correlation curves were fitted with

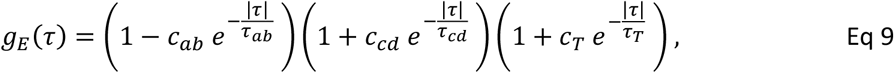

The three terms in the numerator with amplitudes *c*_*ab*_, *c*_*cd*_, *c*_*T*_, and timescales *τ*_*ab*_, *τ*_*cd*_, *τ*_*T*_ describe photon antibunching, chain dynamics, and triplet blinking, respectively. For error estimates of the FRET efficiency correlations, the data were bootstrapped by sampling 1000 times with replacement, and the resulting data sets were fitted with Eq. 9. The total error we report is the root mean square of the standard deviation from bootstrapping and the experimental standard deviation (from replicates if available or, if not, using the average standard deviation).

### Chain reconfiguration time *τ*_*r*_

For any distance-dependent observable, *f*(*r*), the correlation time, *τ*_*f*_, is defined as

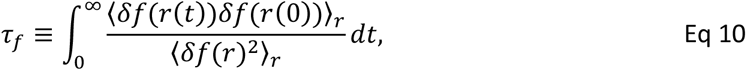

where *δf*(*r*) = *f*(*r*) − ⟨*f*(*r*)⟩_*r*_, and ⟨•⟩_*r*_ = ∫ • *P*(*r*)*dr*. The numerator is defined using the joint probability, *P*(*r*_0_, *r*_*t*_), of populating at an arbitrary time zero the distance *r*_0_ and at a later *t* time the distance *r*_*t*_. With these definitions, we have ⟨*δf*(*r*(*t*)) *δf*(*r*(0))⟩_*r*_ = ∫ ∫ *δf*(*r*_*t*_) *δf*(*r*_0_)*P*(*r*_0_, *r*_*t*_)*dr*_0_*dr*_*t*_. If the dynamics of *r*(*t*) are well described as diffusive motion in a potential of mean force, *F*(*r*) =−*k*_*B*_*T*ln(*r*), then *τ*_*f*_ can be calculated from^124^

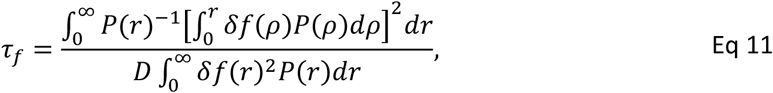

where *D* is the effective end-to-end diffusion coefficient. From fitting the nsFCS curves, we get the intensity correlation time, *τ*_*cd*_ = *τ*_*ϵ*_, where *f*(*r*) = *∈* (*r*) is the transfer efficiency. We can use Eq. 11 to convert to the physically more interesting chain reconfiguration time. We calculated conversion ratios *θ* = *τ*_*cd*_/*τ*_*r*_ for all distance distributions used. *θ* as a function of *R*/*R*_0_ was calculated for the SAW-*v* distance distributions, as well as for the MD simulation ensembles. The difference between simulation and polymer model was propagated as error. Note that *θ* is independent of *D*.

### Polarization-resolved fluorescence lifetime measurements

Lifetime decays from single-molecule experiments were fitted with

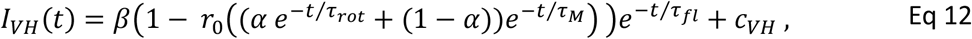

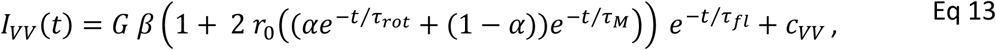

convolved with the instrument response function measured with scattered light. *r*_0_ = 0.38 is the limiting anisotropy of the dyes^125^; *G* accounts for the different detection efficiencies of vertically and horizontally polarized light and was obtained for the donor and acceptor intensities from the ratio of the vertical and horizontal emission after horizontal excitation, *G* = *I*_*HV*_/*I*_*HH*_. The offsets *c*_*VV*_ and *c*_*VH*_ account for background signal. The two rotational correlation times, *τ*_*rot*_ and *τ*_*M*_, account for fast fluorophore rotation and slower tumbling of the entire labeled molecule, respectively. *α* represents the fractional amplitude of the fast component; *β* and *τ*_*fl*_ represent the amplitude and relaxation time of the fluorescence lifetime, respectively.

### NMR spectroscopy

NMR spectra were recorded either on a Bruker Avance III HD 750 MHz spectrometer with a 5 mm TCI Cryoprobe or on a Bruker Avance Neo 800 spectrometer 5 mm CPTXO Cryoprobe. The protein concentration was 0.4 mM for KE_low_ and 1 mM for KE_high_, and for ProTα and the QQEDQQ peptide (TACG Copenhagen, DK) 50 μM and 1.7 mM. For KE_low_ and KE_high_ the buffer composition was 10 mM Tris (pH 6.5), 93 mM KCl, 10 mM DTT, 5% (v/v) D_2_O, including 125 μM sodium trimethylsilylpropanesulfonate (DSS), and the temperature was 25°C or 15°C. For ProTα, the buffer composition was 50 mM Tris (pH 7.4), 0 to 1 M NaCl, 10% D_2_O, 250 µM DSS, and the temperature was 10°C. The 3D spectra of KE_low_ (HNCACB, HNcoCACB, HNCO, HNcaCO, hNcaNH and hNcocaNH) were recorded with 25% nonuniform sampling and processed using qMDD v3.2 using compressed sensing^126^ (ref). Backbone assignments were done manually in CcpNMR Analysis v2.5.2.^126,127^ NMR spectra to investigate salt effects were recorded including ^1^H,^15^N-HSQCs and C_CON spectra.

### CD spectroscopy

Unlabeled protein variants were dialyzed against 10 mM KH_2_PO_4_/K_2_HPO_4_, 10 mM KCl, 1 mM DTT, pH 7.4, using Slide-A-Lyzer MINI Dialysis Devices, 3.5K MWCO (Thermo Fisher Scientific). Insoluble components were removed by centrifugation. Far-UV circular dichroism spectra from 250 nm to 190 nm were acquired on a spectropolarimeter (ChiraScan V100, Applied Photophysics) at 22 °C in quartz cells with a path length of 0.5 mm at concentrations of 0.1–0.5 mg/mL, using an integration time of 0.25 s, averaging 3 measurements, and subtracting the buffer spectrum recorded at identical settings. CD spectra of ProTα were acquired from 260 to 190 nm using a Jasco J-815 spectropolarimeter at 20 °C, with an integration time of 2 seconds. The spectra were averaged over 8 accumulations, and the buffer spectrum, recorded at identical settings, was subtracted. ProTα was measured in a quartz cuvette with a path length of 1 mm, at a protein concentration of 0.1 mg/ml in 10 mM NaH_2_PO_4_/Na_2_HPO_4_ (pH 7.4) buffer, with NaF concentrations between 0 and 500 mM. Absorption data of those scans were used to estimate the concentration of the peptides using their absorption at 214 nm.^128^ Owing to the absence of aromatic residues, the concentration determination has an inherently large uncertainty.

### Quantifying cooperativity

We use cooperativity in an operational sense as the sharpness and collectivity of the response of the conformational ensemble to changes in salt concentration and relate it to equilibrium fluctuations or covariances in the simulated ensembles. As a conceptual starting point rooted in linear response theory, consider an order parameter *m* coupled linearly to an external field *h* that results in a perturbation of the Hamiltonian *H*_0_ as *H* = *H*_0_ − *hm*. In the static equilibrium limit, the corresponding isothermal susceptibility^129^ of the ensemble average of *m*, ⟨*m*⟩, to *h* is

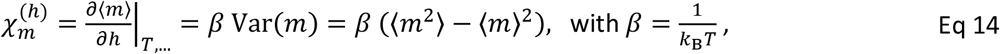

linking a strong response to large equilibrium fluctuations^89^, i.e., a broad free energy surface along *m*. For a cooperative transition, χ_*m*_ typically exhibits a peak near the transition midpoint and provides a measure of cooperativity independent of a two-state assumption^87^ (but χ_*m*_ can of course also be calculated for a two-state system, which we use for comparison). This relation, however, does not directly apply to salt concentration-dependent chain compaction, because ionic strength does not couple linearly to observables such as the end-to-end distance or the FRET efficiency. Instead, in our coarse-grained model, salt affects electrostatic interactions through the Debye-Hückel screening parameter, *k*_D_, with

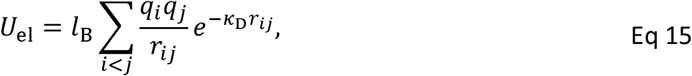

where *l*_*B*_ is the Bjerrum length, and *q*_*i*_ and *q*_j_ are the charges of groups *i* and j separated by distance *r*_*ij*_. The generalized force conjugate to *h*_D_ is therefore

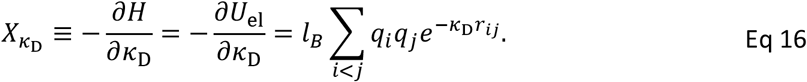

For any observable *A* without explicit dependence on *k*_*D*_, equilibrium statistical mechanics yields the response function or generalized susceptibility^82^

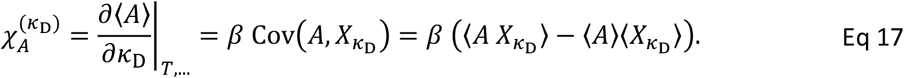

The salt-induced response is thus large if *A* is strongly correlated with the generalized force associated with charge screening. For monovalent salt, 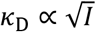, where *I* is the ionic strength, so for the observable as a function of *log*_10_ *I*, as we use it for our analysis,

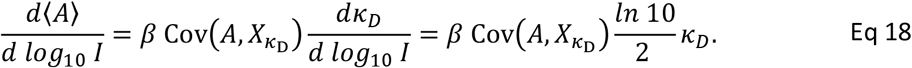

Since 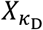 is an energy-like generalized force rather than a directly interpretable structural coordinate, we additionally use the structural surrogate *C*_*net*_ = *C*_*like*_ − *C*_+−_, which tracks the balance of repulsive and attractive electrostatic contacts, *C*_*like*_ and *C*_+−_, respectively. The suitability of a given observable *A* for monitoring the transition can be assessed from its covariance with 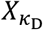 or the linear coupling coefficient 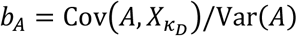. For KE_high_, for instance, 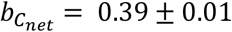and 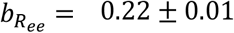 at the midpoint, and correspondingly, Var(*C*_*net*_) is strongly peaked, whereas Var(*R*_*ee*_) is dominated by the expanded ensemble and less peaked (Fig. S4), while the variances of *C*_*net*_ are similar for the compact and expanded ensembles.

From the transitions of observables as a function of ionic strength, ⟨*A*⟩/*dlog*_10_*I* at the midpoint was calculated analytically from the four-parameter sigmoid fit as

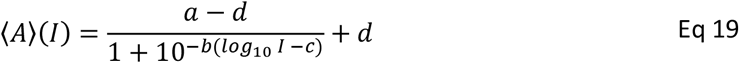

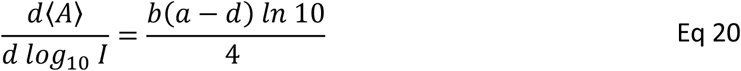

### All-atom explicit-solvent molecular dynamics simulations

Extended all-atom configurations of all six protein variants with explicit Alexa 488 and Alexa 594 dyes were generated using CHARMM^130^ as described previously^131^. The resulting structures were then placed in 14.5-nm rhombic dodecahedron boxes. To prevent self-interaction across periodic boundaries, short vacuum simulations were carried out for each protein, and conformations with a maximum end-to-end distance below 13 nm were retained. Finally, for each of the six variants, three of these conformations were randomly chosen to initiate all-atom explicit-solvent simulations at different salt concentrations. The selected structures were energy-minimized using the steepest-descent algorithm. Each simulation box was then solvated with TIP4P2005s water^60^, and the systems were again energy-minimized using the steepest-descent algorithm. Subsequently, the desired number of potassium and chloride ions were inserted to obtain the following salt concentrations: 10 mM, 400 mM, 500 mM and 1000 mM for KE_high_; 10 mM, 200 mM, 300 mM and 500 mM for KE_mid_; 10 mM, 100 mM, 200 mM and 500 mM for the low charge segregation sequences. Subsequently, the system was again energy-minimized. All simulations were performed using GROMACS^132^ version 2021.5. Protein interactions were modeled using Amber99SBws^60^, with protein-dye and dye-dye interaction parameters described previously^67^ and using optimized dye-water interaction parameters^68^. The temperature was kept constant at 295.15 K using velocity rescaling^133^ (*τ* = 1 ps), and the pressure was kept at 1 bar using the Parrinello-Rahman barostat^134^ (*τ* = 5 ps). Long-range electrostatic interactions were modeled using the particle-mesh Ewald method^135^. Dispersion interactions and short-range repulsion were described by a Lennard-Jones potential with a cutoff at 1 nm. H-bond lengths were constrained using the LINCS algorithm^136^. Each independent run was at least 1.1 μs, and the first 100 ns were treated as equilibration and were omitted from the analysis. For KD_low_ at 10 mM salt, a total of 10 runs were performed and analyzed to ensure sufficient sampling.

### Analysis of MD simulations

Intrachain distance maps between the Cα atoms of each amino acid pair were computed using the distance function of GROMACS^132^ with a saving frequency of 20 ps. Only pairs separated by 3 or more amino acids were used. Correlation times were estimated by integrating the residue Cα-Cα distance autocorrelations, *C*(*t*) (normalized to *C*(0) = 1), up to the time where *C*(*t*) = 0.2 and assuming the remaining decay to be single-exponential^137^. The reconfiguration time of the chain was obtained as the correlation time between the first and the last residue. To obtain the potentials of mean force (PMFs), Boltzmann inversion of the dye-to-dye distance distributions was performed using the C4 atom of the pyran ring of each dye. PMFs from principal component analysis (PCA) and autoencoder models were based on Cα distance maps. Each map was transformed into a one-dimensional array. Subsequently, the arrays were grouped according to salt conditions. For PCA, contact maps were flattened into a one-dimensional array of length n=1,981. 10% of the contact maps from each MD run were randomly selected. Each array was included in a matrix ***X*** of size *m* × *n* with *m*=329,822. The mean *μ* was then computed as

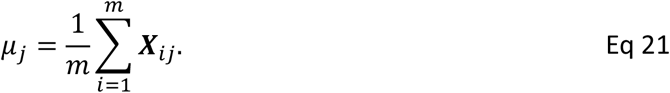

The matrix ***B*** was then calculated as

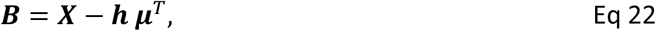

where *h* is a 329,822 x 1 column vector with a constant value of 1. The covariance matrix C was computed as

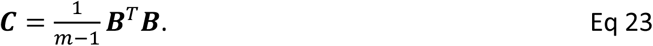

Eigenvectors and eigenvalues of *C* were computed using Mathematica^138^. The eigenvectors were reordered as a function of decreasing eigenvalue. The first two components were employed to project the data into the new space, and the probability distribution was subsequently computed. Finally, Boltzmann inversion was employed on the probability distribution to obtain the potential of mean force. The first two components yielded a fraction of variance explained by the first two principal components of *ρ*^2^ = 0.6 using

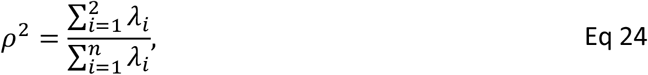

where λ_*i*_ is the i-th eigenvalue of C.

The autoencoder was implemented using Mathematica functions^138^. The neural network contains an input layer (1981×1), a linear layer (657×1), a batch normalization layer, an activation layer, a linear layer (2×1), an activation layer, a linear layer (637×1), a batch normalization layer, an activation layer, and an output layer (1981×1). For the activation layer, a rectified linear unit was used as an activation function. The training process involved the utilization of the ADAM optimizer^138^, with a total of 100 epochs being executed. The model was trained on a set of 80,000 flattened contact maps, of which 20% were used for validation. A dataset comprising 20,000 contact maps was used to assess the performance of the autoencoder after training. A latent space of 2 was found to be the optimal configuration, yielding a coefficient of determination of *ρ*^2^ = 0.82 between the input and output layer at the minimal dimension. Subsequently, the data were projected onto the latent space, and Boltzmann inversion was employed to obtain the PMF from the probability distribution.

The mean transfer efficiency, ⟨*E*⟩, was calculated from MD trajectories as follows: In a first step, the survival probability of the donor excited state with a fluctuating transfer rate, averaged over all possible time origins *t*_0_ along the simulation trajectory, was calculated^67^:

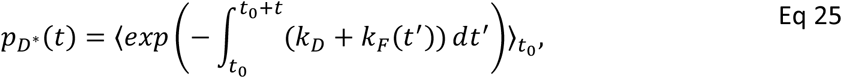

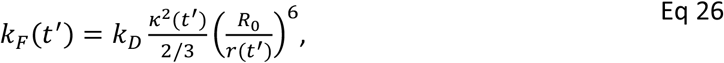

*κ*^2^ denotes the orientational factor for a specific relative orientation of donor and acceptor dye, and *r* the inter-dye distance. *R*_0_ is the Förster radius for *κ*^2^=2/3. *κ*^2^ and *r* were calculated from the MD trajectories using simulation snapshots spaced by 20 ps. The dyes were treated as non-emissive when the distance between any two atoms of donor and acceptor was less than 0.4 nm, corresponding to van-der-Waals contact^139,140^. The average FRET efficiency, ⟨*E*⟩, was calculated by integration over 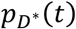:_67,141_

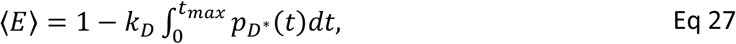

where *t*_*max*_ = 20 ns represents the time after excitation by which 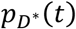 is essentially zero^67^.

Lifetimes and fraction bound of residue–residue contacts were calculated by a transition-based or core-state approach^142^. In short, rather than using a single distance cutoff to separate bound versus unbound states, which tends to underestimate contact lifetimes, separate cutoffs were used to determine the formation and breaking of contacts. For each pair of residues, a contact was based on the shortest distance between any pair of heavy atoms, one from each residue. Starting from an unformed contact, contact formation was defined to occur when this distance dropped below 0.38 nm; an existing contact was considered to remain formed until the distance increased to more than 0.8 nm.^142^

Dynamics of the interchromophore distance was analyzed with a one-dimensional diffusion model^73,104,143^. Briefly, the distance coordinate was discretized into 30 equal-size, nonoverlapping bins between the minimum and maximum sampled value for each trajectory. Histograms of the number of observed transitions between bin *i* at time *t* and bin *j* at time *t* + Δ*t, N*_*ij*_ (Δ*t*), were determined for different lag times Δ*t*. Discretized free energies and position-dependent diffusion coefficients were optimized via Monte Carlo simulations using the log-likelihood function.

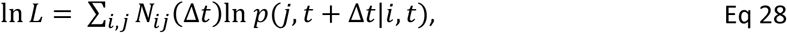

where the propagators *p*(*j, t* + Δ*t*|*i, t*) describing the conditional probability of being in bin *j* a time Δ*t* after having been in bin *i* are obtained from the discretized diffusion model as previously described^104,143^: in short, the discretized dynamics is mapped to a chemical kinetics scheme describing the time evolution of populations in the bins, 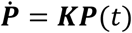 where ***P***(*t*) is the vector of the bin populations at time *t*, and ***K*** is a rate matrix derived from the diffusion coefficient(s) *D*_*i*_and free energies *F*_*i*_ associated with each bin according to the scheme of Bicout and Szabo^143,144^. The propagators are then given by *p*(*j, t* + Δ*t*|*i, t*) = (*exp*(Δ*t****K***))_j*i*_ . In estimating the most probable parameters from the data, a uniform prior is used for the diffusivities and free energies. We can compute normalized correlation functions.

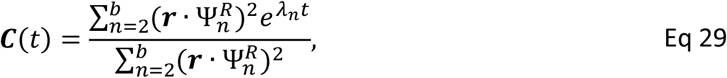

where the elements of ***r*** are the centers of each bin on the distance coordinate, 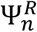is the *n*th right eigenvector of ***K*(** 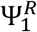 is the stationary eigenvector), and λ_*n*_ is the *n*th eigenvalue. From this, the correlation times follow:

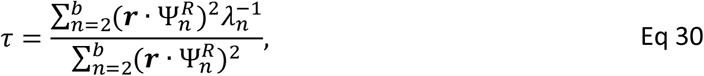

In determining the optimal parameters, we fitted simultaneously to lag times of 5, 10, and 20 ns, so as to include both short- and long-time correlations.

### Coarse-grained molecular dynamics simulations

Proteins were modeled at a single-bead-per-amino-acid resolution, with each bead representing an amino acid centered at its Cα position, and the fluorescent dye pairs (Alexa 488 and Alexa 594) were explicitly represented as described before^31^. The beads are connected using a harmonic bond potential with a force constant of 418 kJ nm^-2^ mol^-1^ and an equilibrium bond length of 0.38 nm. Electrostatic interactions were modeled using a screened Coulomb potential, and the non-electrostatic components were modeled using a short-range solvation potential (hydrophobic scale/HPS model) that included the parameters for explicit dyes^31^. The proteins were placed in the center of a 17×17×17 nm box with periodic boundary conditions. Langevin dynamics simulations were performed using GROMACS^132^ at 300 K with a time step of 10 fs and a friction coefficient γ of 1 ps^-1^. The simulations were performed for 1.5×10^8^ MD steps; the first 5×10^7^ steps were discarded as equilibration, and coordinates were stored every 10000 steps.

Mean FRET efficiencies were calculated from the simulations using the inter-dye distances using the Förster equation:

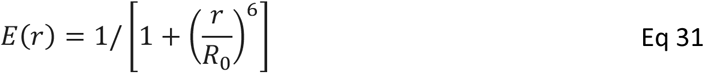

Here, *r* is the distance between the dye beads^31^, and ***R***_*o*_= 5.4 nm is the Förster radius of the Alexa dye pair used in the experiments.The ⟨*E*⟩ values are compared with the experimental FRET efficiencies. The trajectories were analyzed in Python using the MDAnalysis library^145^, and intramolecular contacts were calculated using a 1.2 nm distance cutoff, excluding the dye beads.

The slope at the midpoint of the ionic strength-dependent transitions was calculated from Eqs. 19 and 20. The KE_mix_ and KE_seg_ sequence libraries (128 protein sequences each) were generated by randomizing the KE_high_ sequences, treating KE as a unit, or by segregating charged residues at the termini, respectively. The hypothetical two-state reference for KE_seg_ was estimated by treating the conformational ensembles at 8 mM (compact; c) and 1008 mM (expanded; e) as fixed states and changing only their relative populations with salt concentration. The free-energy difference between the two states as a function of salt concentration was calculated as

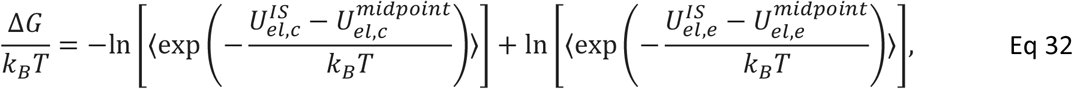

where 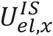 is the electrostatic interaction energy of a configuration at a given ionic strength (IS), 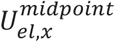 is the electrostatic interaction energy of the same configuration at the midpoint salt concentration, and averaging is over all configurations of the ensemble. The probabilities of the compact (*p*_***c***_) and expanded (*p*_***e***_) states were calculated from

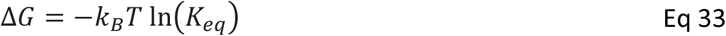

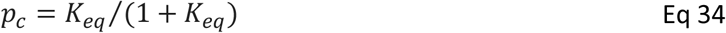

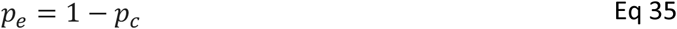

The mean value for any observable or reaction coordinate *A* of the two-state reference is

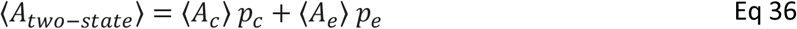

and the variance as

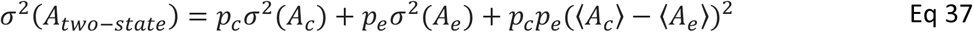

The two-state free-energy profiles were constructed by summing up the probability distributions of the observables at 8 and 1008 mM ionic strength. PyMOL^146^ was used for molecular visualizations.

### Sequence Charge Decoration

Human proteome-wide intrinsically disordered region (IDR) sequences were obtained from Tesei *et al*.^12^ and used to calculate the sequence charge decoration (SCD), defined as previously described^25^:

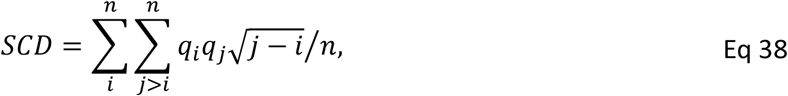

where *q*_*i*_ and *q*_*j*_ are the charges of residues *i* and *j*, respectively, and *n* is the sequence length. Aspartate (D) and glutamate (E) residues were assigned a charge of −1, while lysine (K) and arginine (R) residues were assigned a charge of +1. Only the negative arm of the SCD distribution is shown in Figure 1d, as this regime includes polyampholyte sequences satisfying the following criteria: (a) sequence length ≤150 amino acids, (b) fraction of charged residues (K, R, D, and E) >30%, and (c) absolute net charge ≤5% of the sequence length. These criteria match those of the sequences studied in this work.

## Acknowledgements

We thank Nicola Galvanetto and Andrea Sottini for help with data analysis and helpful discussions, Marie Synakewicz for help with plasmid design, Freia S. Buus for recording the QQEDQQ data, Milena Srejic for help with some of the single-molecule measurements, and Hagen Hofmann, Attila Szabo, and Wenwei Zheng for insightful comments on the manuscript. We thank the Functional Genomics Center Zurich for mass spectrometry analysis. This work was supported by the Swiss National Science Foundation (to B.S., 310030_197776 and CRSII5_205922), the Forschungskredit of the University of Zurich (00109396 to S.G.) and the Novo Nordisk Foundation Challenge grant REPIN – rethinking protein interactions (#NNF18OC0033926 to B.B.K. and B.S.). R.B.B. was supported by the Intramural Research Program of the National Institute of Diabetes and Digestive and Kidney Diseases of the National Institutes of Health. We used the computational resources of Piz Daint, Alps, and Eiger at the CSCS Swiss National Supercomputing Center, and of the National Institutes of Health HPC Biowulf cluster (http://hpc.nih.gov). We thank the Novo Nordisk Foundation and Villum Fonden for supporting the NMR facility at UCPH, DK.

