## Supplementary Information for "Cooperativity, dynamics, and the free-energy surfaces of charge-patterned IDPs"

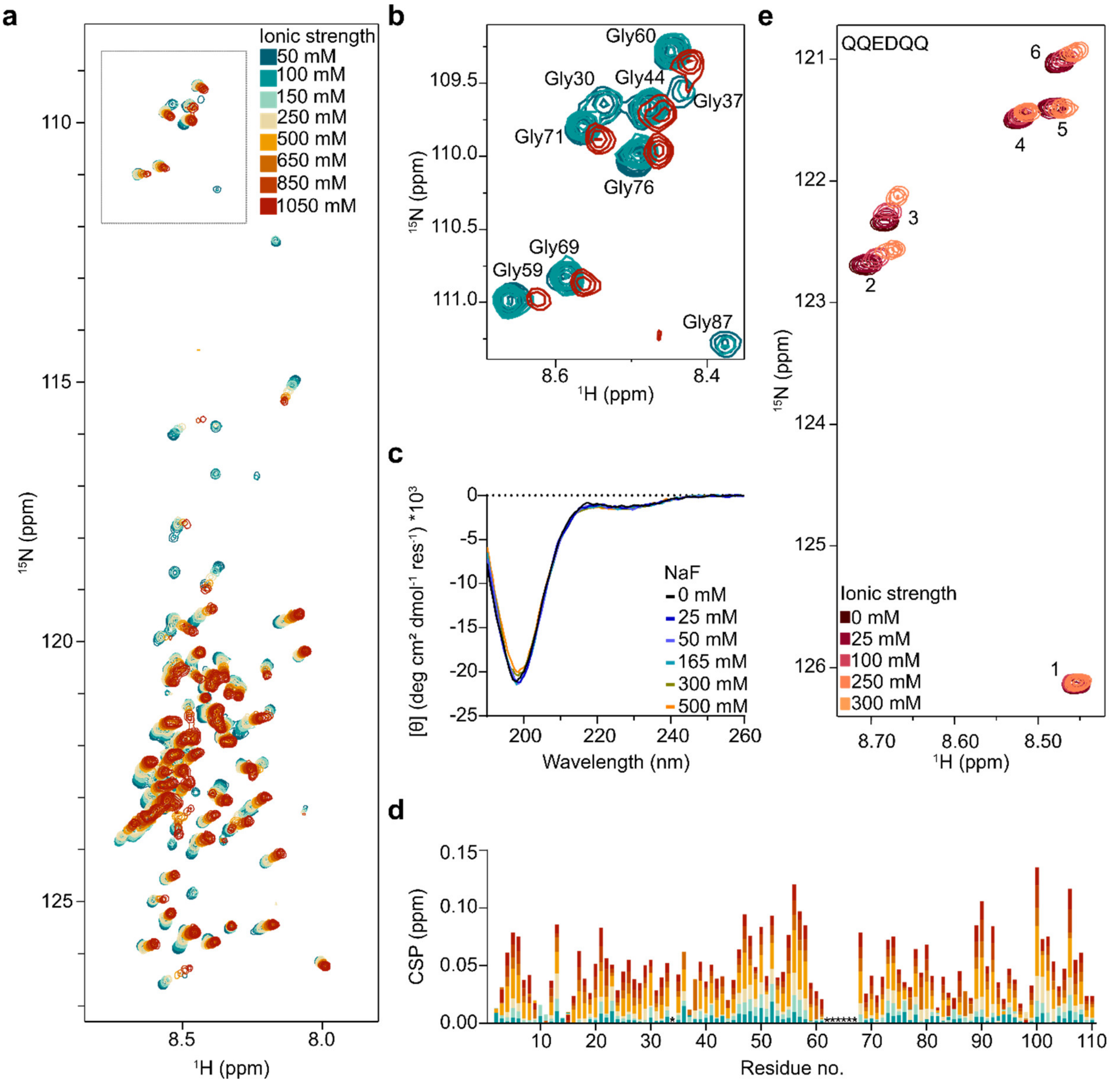


***Figure S1: Increasing salt concentration does not induce secondary or tertiary structure in ProTα and a random coil peptide.*** *(**a**) $^{1}$H,$^{15}$N-HSQC spectra of ProTα* at *different NaCl concentrations. (**b**)* Zoomed-in region of the $^{1}$H-$^{15}$N-HSQC spectra of $^{15}$N-ProTα *at different NaCl concentrations (50, 100, 1000 mM). Asterisks correspond to unassigned residues. (**c**) Far-UV CD spectra of ProTα at different NaF concentrations confirm the absence of changes in secondary structure content. (**d**) Chemical shift perturbations (CSP) of $^{15}$N-ProTα at different NaCl concentrations. (**e**) $^{1}$H,$^{15}$N-HSQC spectra of QQEDQQ peptide recorded at natural abundance at different KCl concentrations. Peaks 2 and 3 correspond to the Glu and Asp residues in the peptide (unassigned), illustrating the generic changes in chemical shifts with increasing salt concentration.*

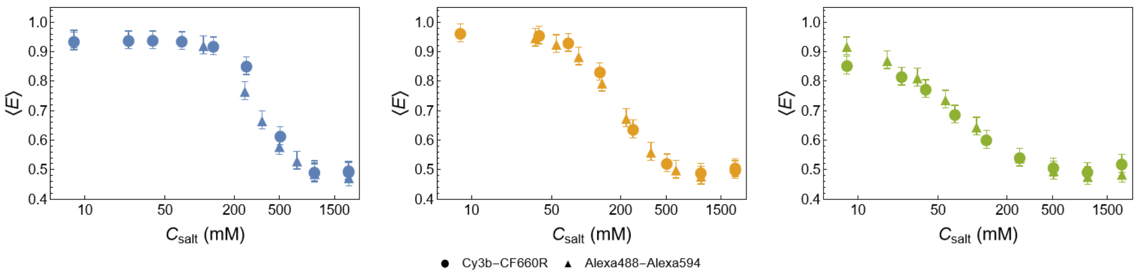


***Figure S2: FRET labels have no strong effect on salt concentration-dependent chain expansion.*** *Comparison of experimental single-molecule transfer efficiency as a function of salt concentration for* $KE_{high}$ *(left),* $KE_{mid}$ *(middle) and* $KE_{low}$ *(right) with the dye pairs Alexa Fluors 488/594 and Cy3B/CF660R (see legend).*

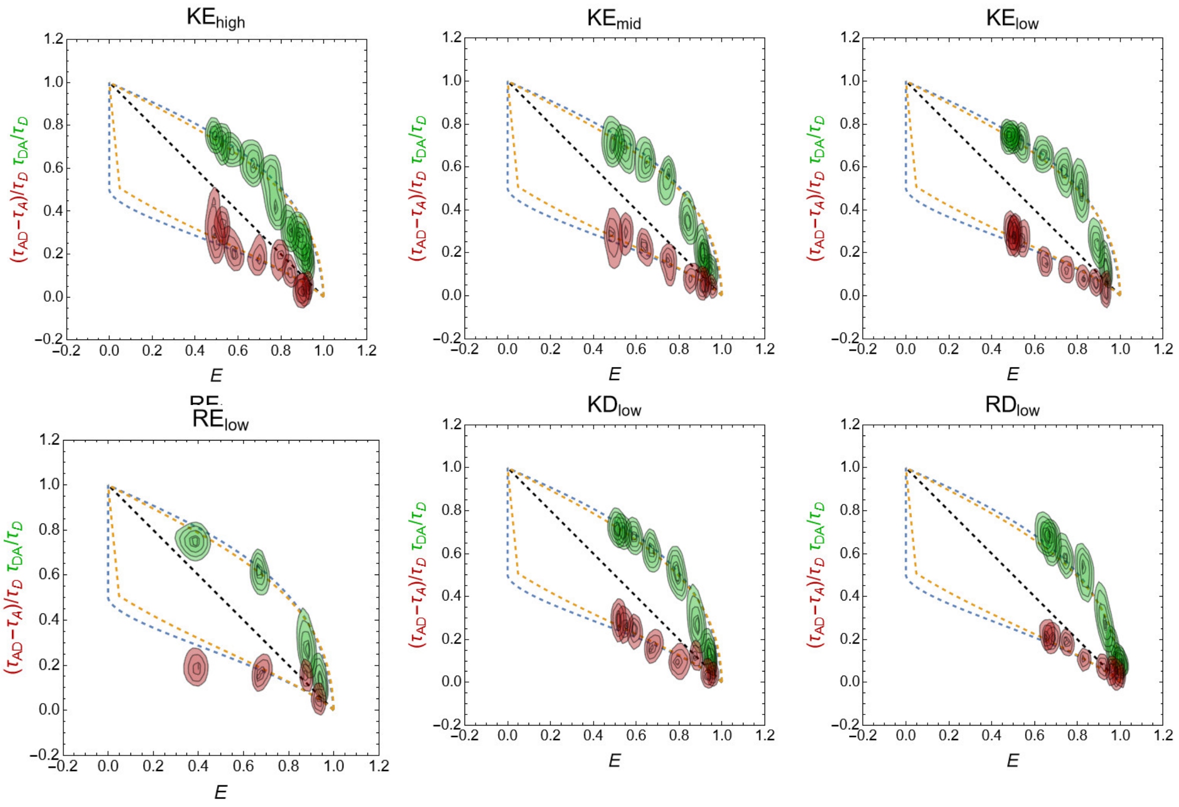


***Figure S3: Transfer efficiency vs fluorescence lifetime for all protein variants from single molecule experiments indicate rapidly sampled broad distance distributions.[1-3]*** *The diagonal black dashed lines indicate the relation expected for static distances. The orange dashed lines show the relation for a rapidly sampled SAWv distance distribution[4], and the blue dashed lines for a Gaussian chain.*

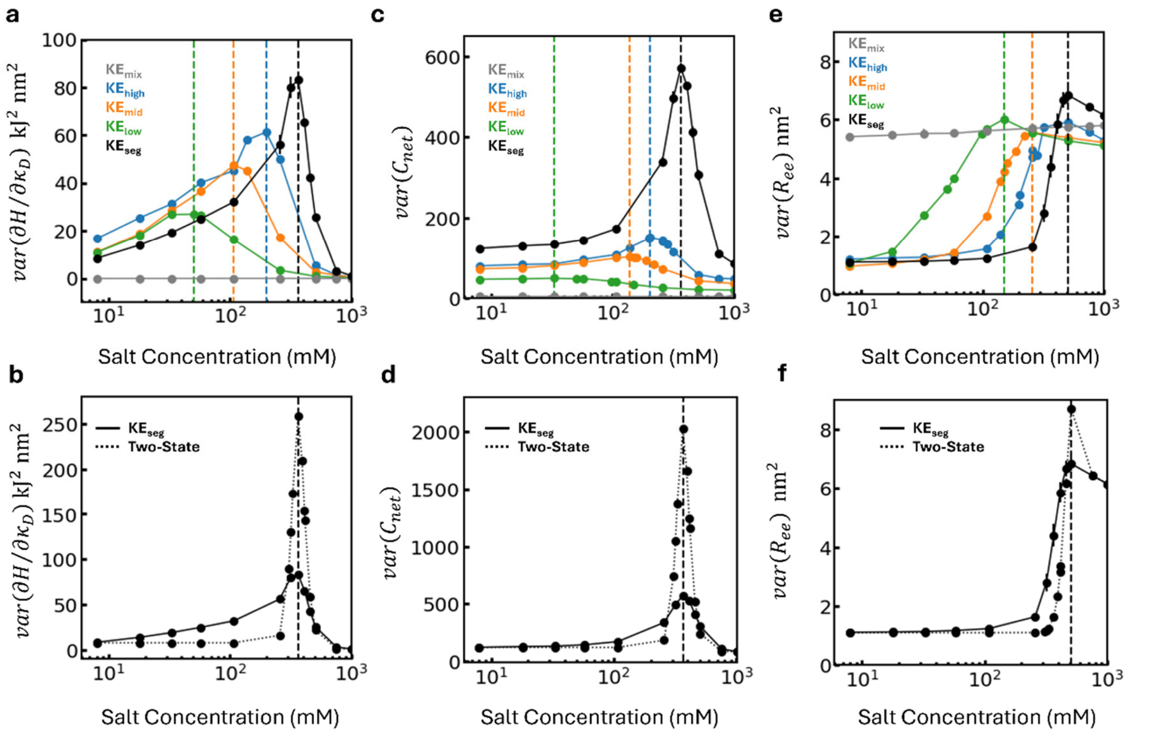


***Figure S4: Comparing the salt concentration-dependent variances for different order parameters.*** *Variance of (a,b)* $\partial H/\partial\kappa_D$*, (c,d) net contact balance (*$C_{net}$*), and (e,f) end-to-end distance (*$R_{ee}$*) of* $KE_{seg}$*,* $KE_{high}$*,* $KE_{mid}$*,* $KE_{low}$*, and* $KE_{mix}$ *as a function of salt concentration from coarse-grained simulations. The variance of two-state references are calculated using Eq. 37. The vertical dashed lines mark the inflection points of the transitions.*

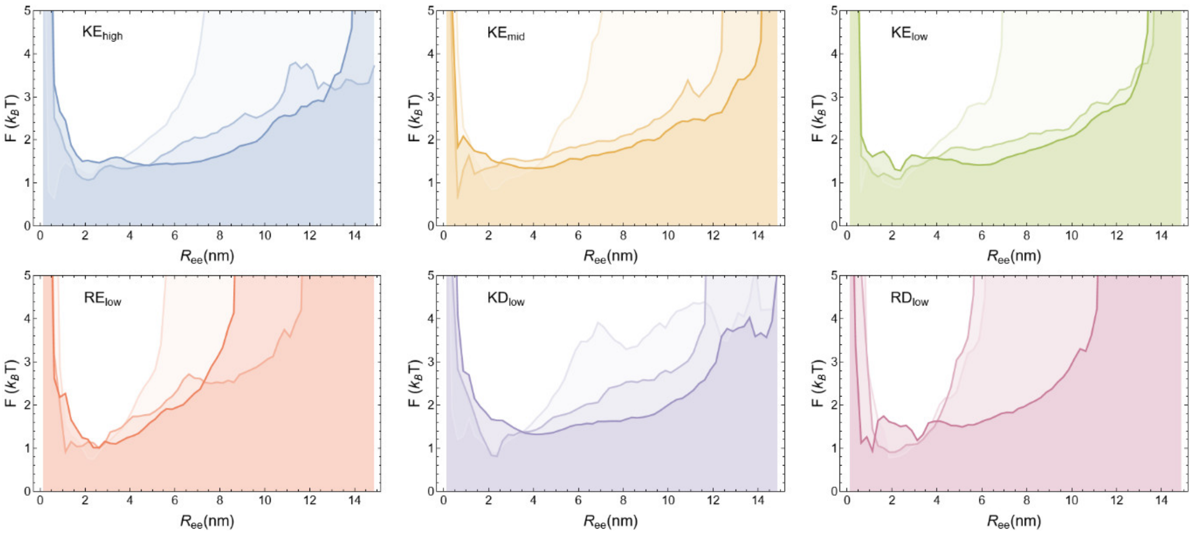


***Figure S5: Potentials of mean force for all protein variants from Boltzmann inversion of the dye-dye distances, $R_{ee}$, from all-atom simulations do not show pronounced barriers.*** *Color saturation indicates salt concentration. Lighter color represents lower salt concentration. Low salt is 10 mM for all variants, medium salt is 400 mM for $KE_{high}$, 200 mM for $KE_{mid}$, and 100 mM for the others. High salt is 1000 mM for $KE_{high}$ and 500 mM for the others.*

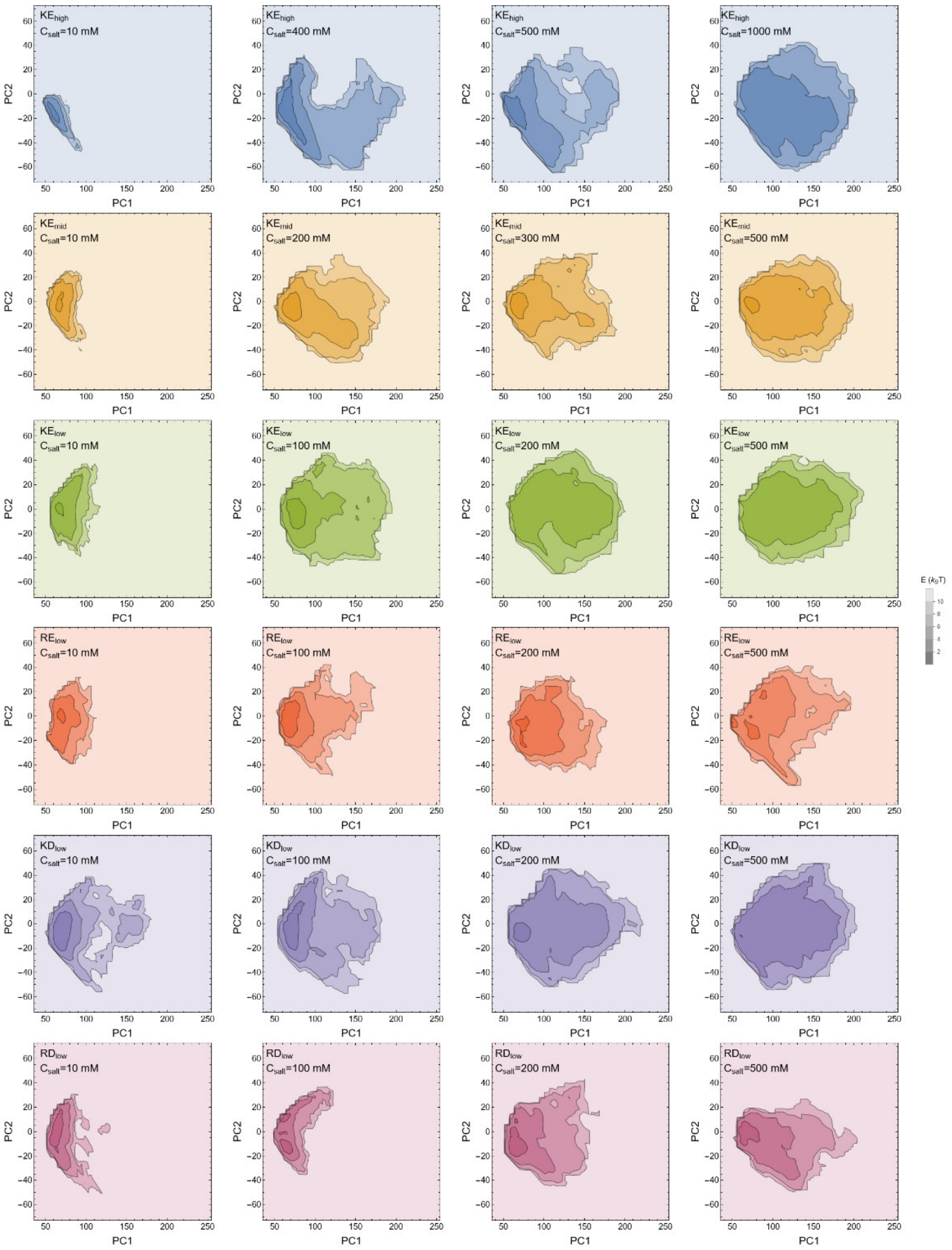


***Figure S6: Principal component analysis does not identify pronounced barriers.*** *Projection of the free energy from the conformational distributions of all protein variants based on all-atom MD simulations on the dimensions from principal component analysis. Color saturation indicates free energy (see scale).*

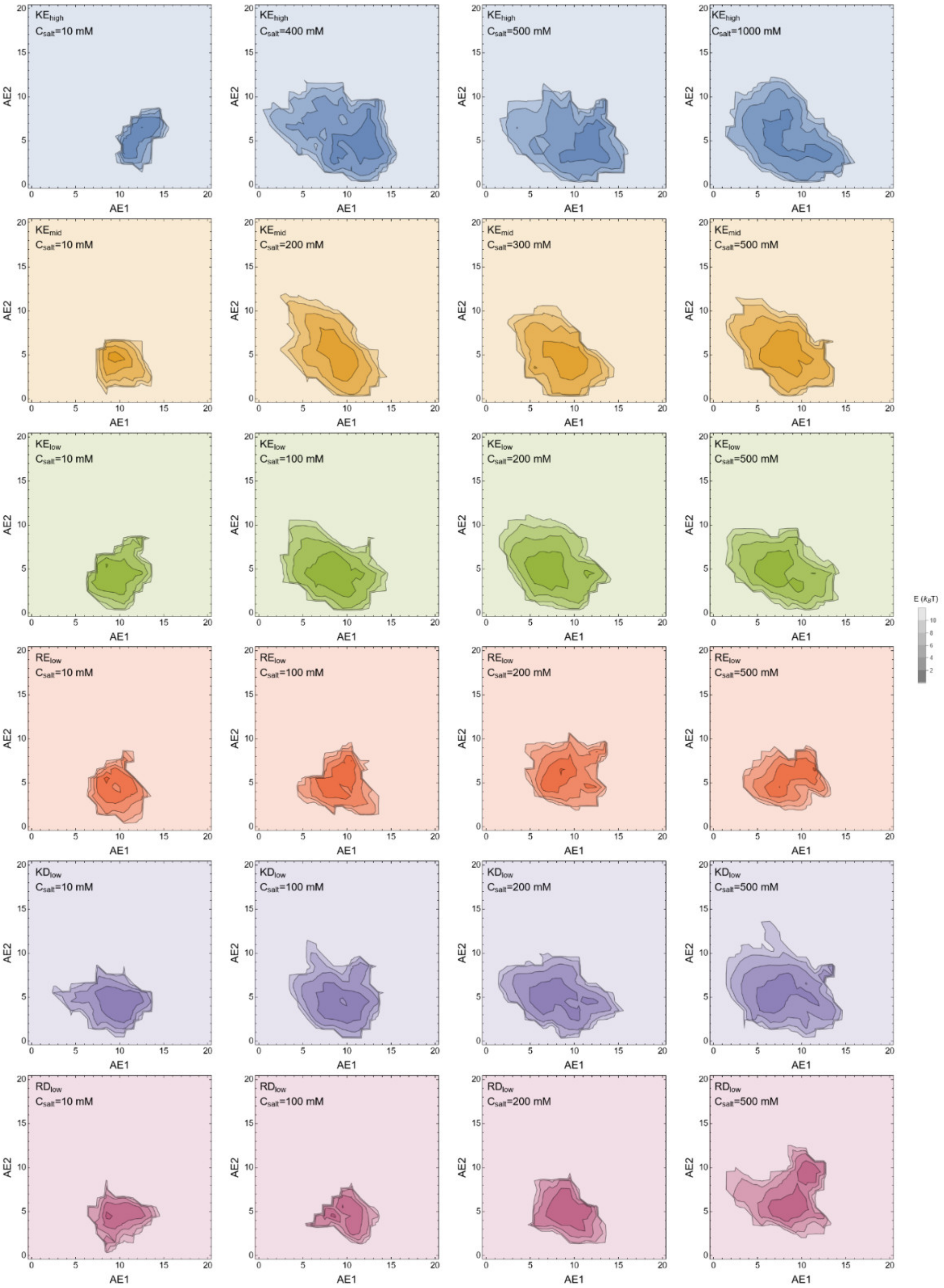


***Figure S7: Autoencoder analysis does not identify pronounced barriers.*** *Projection of the free energy from the conformational distributions of all protein variants based on all-atom MD simulations on the dimensions from the autoencoder analysis. Color saturation indicates free energy (see scale).*

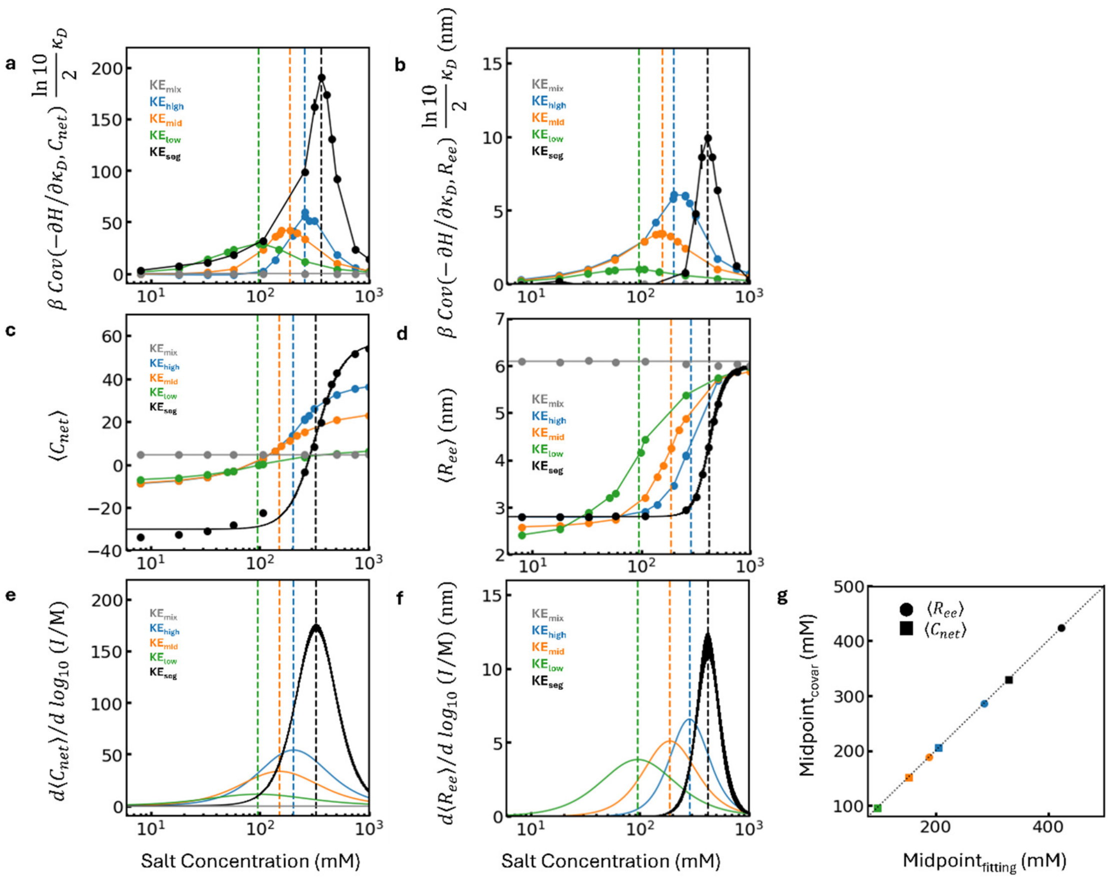


***Figure S8: Susceptibility as a measure of cooperativity.*** *(a,b) Susceptibility calculated from $\beta\; Cov\left(A, X_{\kappa_D}\right)\frac{ln\,10}{2}\kappa_D$ (Eq. 18), where A is $C_{net}$ or $R_{ee}$ for KE$_{seg}$, KE$_{high}$, KE$_{mid}$, KE$_{low}$, KE$_{mix}$ as a function of salt concentration from coarse-grained simulations. (c,d) Salt-dependent transitions of $\langle C_{net}\rangle$ or $\langle R_{ee}\rangle$. The solid lines are the fit to the data. (e,f) Susceptibility calculated numerically from the slopes of the sigmoidal fits to salt-dependent transitions, $d\langle A\rangle/d\;log_{10}I$, in panels (c,d). The vertical dashed lines mark the midpoint. (g) Comparison of the midpoints obtained from the sigmoidal fits of the transitions (Eq. 19; panels c,d) and from the maxima of the covariance-based analysis (a,b).*

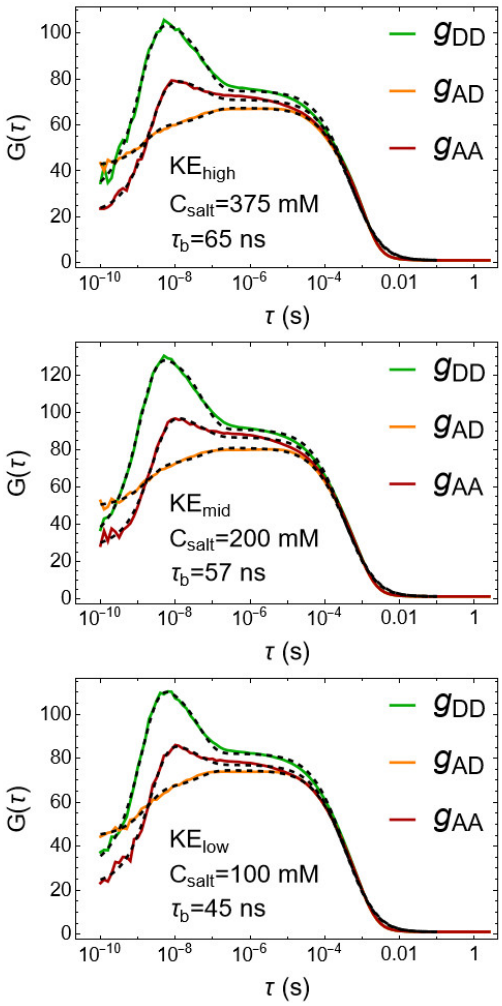


***Figure S9: Fluorescence correlation spectroscopy using samples with Cy3B-CF660R indicates the absence of distance dynamics above 100 ns.*** *KE$_{high}$, KE$_{mid}$ and KE$_{low}$ labeled with Cy3B-CF660R were measured at their midpoint salt concentrations. Dashed lines indicate fits with Eq. 7.*

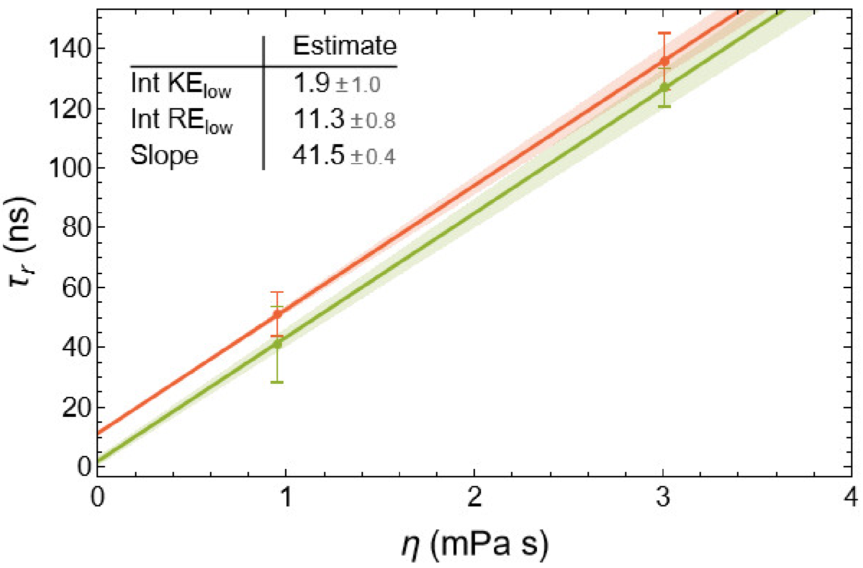


***Figure S10: Small difference in internal friction between KE$_{low}$ and RE$_{low}$ at low salt concentration.*** *Solvent viscosity-dependent reconfiguration times, $\tau_r$, of KE$_{low}$ and RE$_{low}$ from nsFCS at 100 mM salt concentration (lines: linear fits; inset: internal friction times in nanoseconds; error bars from bootstrapping, see Methods).*
